# Genetic heritage of the BaPhuthi highlights an over ethnicised notion of ‘Bushman’ in the Maloti-Drakensberg, Southern Africa

**DOI:** 10.1101/2022.03.05.483114

**Authors:** Ryan Joseph Daniels, Maria Eugenia D’Amato, Mpasi Lesaoana, Mohaimin Kasu, Karen Ehlers, Paballo Abel Chauke, Puseletso Lecheko, Sam Challis, Kirk Rockett, Francesco Montinaro, Miguel González-Santos, Cristian Capelli

## Abstract

Using contemporary people as proxies for ancient communities is a contentious but necessary practice in anthropology. In Southern Africa, the distinction between the Cape KhoeSan and eastern KhoeSan remains unclear as ethnicity labels are continually changed through time and most communities were extirpated. The eastern KhoeSan may reflect an ‘essentialistic’ biological distinction from neighbouring Bantu-speaking communities or it may not be tied to ‘race’ and instead denote communities with a nomadic ‘life-way’ distinct from agro-pastoralism. The BaPhuthi of the 1800s in the Maloti-Drakensberg, Southern Africa had a substantial San constituency and a life-way of nomadism, cattle raiding, and horticulture. The BaPhuthi heritage could provide insights into the history of the eastern KhoeSan. We examine for the first time genetic affinities of 23 BaPhuthi to distinguish if KhoeSan ancestry reflects biologically distinct heritage or a shared life-way. Data were merged with 52 global populations. The Principle Component Analysis, ADMIXTURE clustering and F_3_ tests show no support for a unique eastern KhoeSan ancestry distinct from other KhoeSan or southern Bantu-speaking communities. The BaPhuthi have strong affinities with Nguni communities, as the non-Nguni show strong evidence of recent African admixture possibly related to late-iron age migrations. The BaPhuthi may have an interesting connection to the early iron-age Bantu-speaking communities as MALDER detected no signals for late-iron age admixture. We demonstrate how the ‘essentialistic’ understanding of references in historic literature creates misconstrued notions of ethnic/biological distinctions when ‘San’ and ‘Bushman’ may have reflected ambiguous references to the non-sedentary polities and practices.

## 1 Introduction

Using contemporary people as proxies for ancient communities is a contentious practise in anthropology (1-5) and an ongoing discussion in trying to understand the relationship between KhoeSan peoples and culture in Southern Africa (6-8). As communities and cultures are continually re-invented and lost, only an imperfect account of the past can be gathered from extant cultures and people (2). Researchers look toward physical remains and historic accounts as well but connecting past descriptions to contemporary peoples too may be confounded. Identity labels are continually formed, morphed, appropriated and lost through time.

KhoeSan refers to the collective of linguistically and culturally diverse African communities from a range of environments, regions and times (10; 11). Possible cultural, biological and/or linguistic distinctions between eastern and western KhoeSan are incompletely understood (9). The term ‘KhoeSan’ and the many pre-cursor terms are contentious because their historic use and we refer readers to Text_S 1 for fuller discussion on the terms used here. In the western parts of South Africa, KhoeSan have often been referred to as ‘San’, ‘Hottentots’, or ‘Khoikhoi’ (9; 12) while those who inhabited mountainous regions of eastern Southern Africa - present-day Lesotho, KwaZulu-Natal, Griqualand East, and the former Transkei - have been referred to as ‘Bushman’, ‘Mountain Bushmen’ or ‘People of the Eland’ (9; 13-15). In terms of language and identity, communities have persist in the western regions although with clear influence from historic events. In eastern Southern Africa, however, there are no known remnant KhoeSan communities from which to draw insights.

Any possible ancient signals of divergence are made more complex by historic and ongoing developments. Clear influences are found in cultural diffusion and genetic exchange associated with a number of events. For example, the arrival of the East African pastoralists to Southern Africa ∼2000 years ago introduced Eurasian genes and livestock (6; 11; 16; 17). The subsequent extensive spread of Bantu-speaking communities brought iron-technology, sedentary agro-pastoralism and socio-political change (11; 18-21). The mounting pressure from European colonial expansion and Bantu-speakers’ nation-building during the 1600s – 1900s dissolved several KhoeSan cultures and people entirely. Such ‘vanished’ communities are known largely, if not entirely, from historic texts and often with sporadic, disjointed mentions.

What we know of the western KhoeSan were detailed from early encounters with Europeans which have provided insight into pre-colonial communities (9; 22; 23). The ‘San’ were described as smaller in stature and paler of skin than Southern African Khoikhoi and agro-pastoralists (12; 24). Other identifying features included the tightness of hair curls, eye folds and cranial structure (25-27). Furthermore, linguistic work on contemporary people and from historic accounts have allowed the mapping of possible distributions of linguistically identifiable groups, and relationship between San communities in the West. These details are largely lacking for KhoeSan in the East.

Some information can be gathered from references to ‘Bushman raiders’ in the seminal work of Vinniecomb and Wright (14; 28). Here ‘Bushman’ are described as akin to the San however, the term may be hyper-ethnicised. For example, the division between ‘San’ and non-’San’ is rooted in and perpetuated by colonial tendencies to exaggerated physical differences (29). The connotation of ‘Bushman’ to African and European authorities in the 19th century was a pejorative use of the word rather than an ethnic identity (29). Terms such as ‘San’, ‘BaTwa’, and ‘BaRoa’ may to have instead denoted a ‘life-way’, and not just ‘race’ (8). The work by Rachel King and Sam Challis further argues that one could adopt the life-way and become a ‘Bushman’ and that ‘Bushman’ communities may have been heterogeneous only sharing a life-way which spurned sedentary polities in favour of hunting, gathering and live-stock raiding (8; 29). Indeed, close relations between KhoeSan and Bantu-speaking communities are a characteristic of the Maloti-Drakensberg history (29). Creolised identities such as the AmaTola and BaPhuthi are a testament to this (30).

The contemporary BaPhuthi of the southern Maloti-Drakensberg speak a Nguni Bantu language, SePhuthi, and were reported to be ethnically heterogeneous in the 1800s with a substantial KhoeSan constituency (12; 31; 32). The BaPhuthi of the 1800s rejected the idea of a sedentary ‘Great Place’ chieftaincy in favour of circulating through a series of settlements atop steep sided *kopjes* or *liqhobosheane* scattered along the Senqu river (12). Their life-way was based on nomadism, cattle raiding, and horticulture rather than field-based agriculture; these are traits shared with their KhoeSan antecedents and contemporaries (33). Moorosi, who led the BaPhuthi in the 1800s, had KhoeSan and Bantu-speaking wives (33). The AmaTola of the Eastern Cape reportedly assimilated Khoikhoi pastoralist during the 18^th^ century Frontier Wars (8; 29; 30) and some of their members were subsequently assimilated into the BaPhuthi (33). As with the Lake Chrissie communities who report descent from ||Xegwi speaking San and do indeed have elevated KhoeSan descent (34), the BaPhuthi’s oral history may be reflected in genetic signatures of the eastern KhoeSan.

It remains unclear to what extent the narratives are signs of an assimilation of KhoeSan peoples and culture or rather the adoption of the practises without direct KhoeSan descent. There are thus two contending ideas for the ‘Bushman’ descent of the BaPhuthi; 1. a possible descent from a ‘San’-type nomadic community with some ‘essentialistic distinction’ from the western KhoeSan or 2. that ‘Bushman’ were heterogeneous societies with a shared life-way and may not have had any, or very limited, ‘San’-type biological affinities.

Efforts toward understanding the genetic diversity of Southern African KhoeSan and the history of the region increasingly requires the insights from remnant genetic signals in descendant communities, such as the BaPhuthi. To this end, we examine for the first time the genetic affinities of BaPhuthi individuals with oral history of KhoeSan descent from south of the Maloti-Drakensberg.

## 2 Participants and Methods

### 2.1 Ethics Approval

Ethics approval for the South African samples was obtained from Oxford Tropical Research Ethics Committee (Oxford University, UK) (ref. No. 8-16) and the University of the Free State (South Africa) NatAgri Ethics Committee (ref. No. UFS-HSD2016/1210). Approval for the Lesotho samples was granted by the University of the Western Cape Research Ethics Committee (ref. No. BM16/3/18) and the Ministry of Health, Lesotho (ID128-20016). Export permits were approved by the South African Department of Health and the Lesotho Ministry of Health.

### 2.2 Sample Collection and Genotyping

We conducted interviews with residents from Masakala in South Africa (2017) and from Semonkong village and Quthing district in Lesotho (2019) (Figure 1). Information on mother-tongue language, place of birth and ethnicity of the participant, their parents and grandparents were collected. Approximately 2ml of saliva was collected with Oragene-500 kits (DNA Genotek, Canada). Where participants indicated that all four grandparents were BaPhuthi or spoke SePhuthi, samples were used for genotyping. DNA extractions were performed at Oxford University following the prepIT.L2P salt extraction protocol (catalogue# PT-L2P, DNA Genotek, Ottawa Canada). A total of 33 samples from Lesotho were genotyped for ∼600K SNPs on the Genome Screening Array (GSA v2) at the Estonian Institute for Genomics, Tartu, Estonia. A further two samples from South Africa were genotyped for 2.5 million variants on the Illumina Omni2.5-8 Beadchip v1.3 at the Wellcome Trust Centre for Human Genomics, Oxford University. All raw output was processed using GenomeStudio software (Illumina, USA) and all samples passed a call rate of 97% or more.

**Figure 1:**
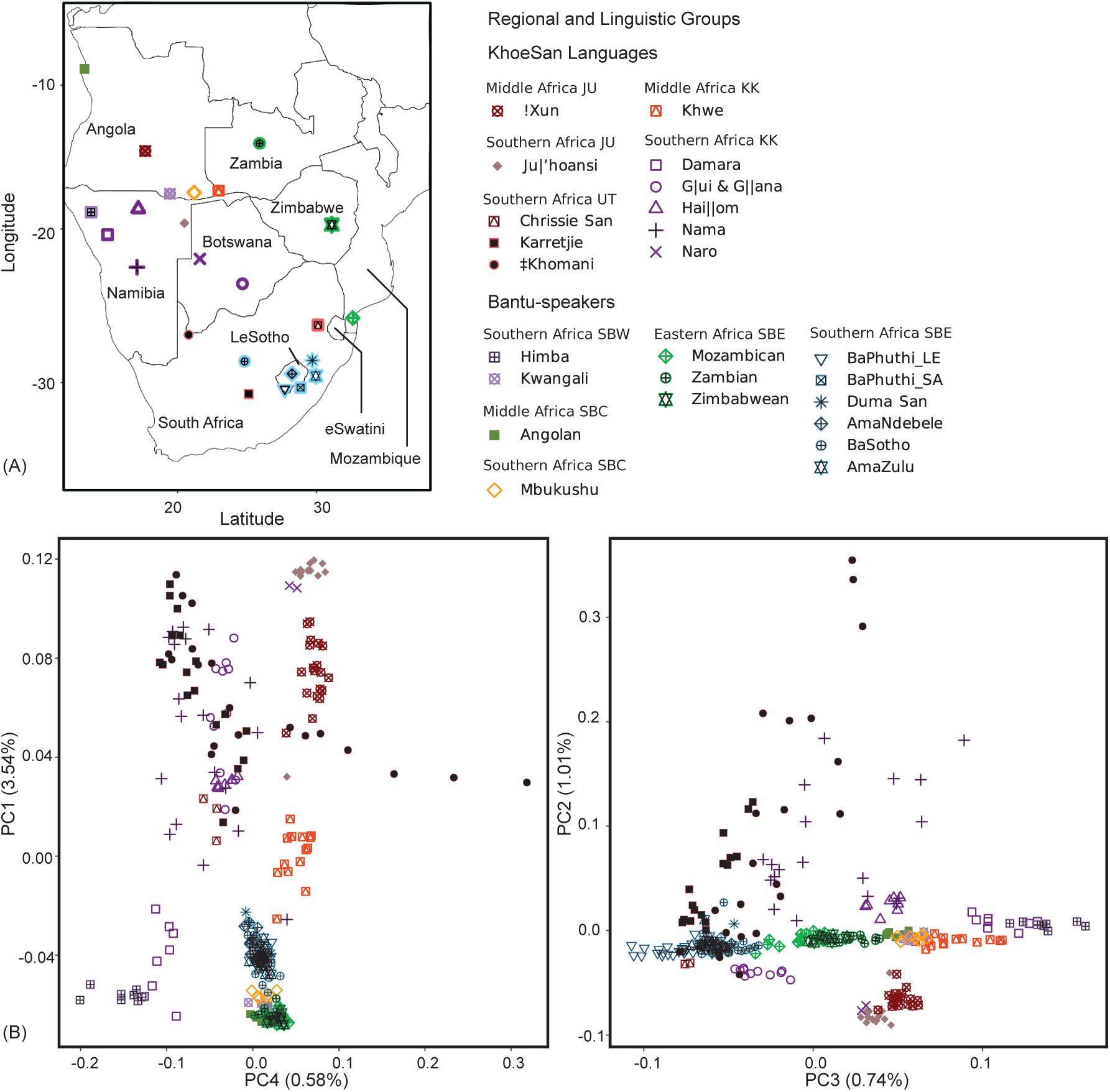
(A) A Map of the Bantu-speaking and KhoeSan populations included in the analyses. Country labels included. (B) Principal component (PC) analysis showing the first four PCs arranged to emphasis patterns. The percentage variation explained by each component is indicated in brackets at the respective axis. Colours indicate regional and linguistic divisions. The focal populations are indicated by a overlain black symbols. Linguistic abbreviations used: southern Bantoid, western Bantu – SBW, southern Bantoid, Central western Bantu – SBC, southern Bantoid, eastern Bantu – SBE, Juu KhoeSan – JU, Khoe-Kwadi KhoeSan – KK, Ui! and Taa KhoeSan – UT.

### 2.3 Datasets, Merging and Quality Control

#### 2.3.1 Data

We merged the samples genotyped here with publicly available data genotyped on the Illumina Omni5 or 2.5 array from the African Genome Variation Project (35), the 1000 Genomes Project (36), and four Southern African datasets (34; 37-39). The latter included a collection of 11 KhoeSan groups including representatives of the Khoi-Kwadi (abbreviated to KK), Juu (JU) and !Ui - Taa (UT) language areas (Figure 1A). In all populations except the BaPhuthi we restricted sample sizes to 20 individuals to reduce computation load.

The final dataset included a collection of Southern African populations to which we pay particular attention to compare with the BaPhuthi. This focal group included several Southern African KhoeSan groups (Karretjie, ‡Khomani and Namibian Nama as the only representative of Khoikhoi groups) and two groups of Bantu-speakers who are geographically and linguistically close to the BaPhuthi (Lake Chrissie San on the border of Eswatini, Duma San from the KwaZulu-Natal uKhahlamba-Drakensberg). We additionally included several representatives of southern Bantoid, eastern Bantu speaking communities (here abbreviated to SBE). These included individuals from Zambia and Zimbabwe (both from Eastern Africa (40)), the Himba of Namibia (southern Bantoid, western Bantu; SBW), and several Southern African SBE (BaSotho, AmaZulu, and AmaNdebele). The remaining samples made up a ‘global reference’ set from which we infer affinities.

From each data set, the following were conducted in PLINK (41). We retain only bi-allelic variants and pruned for T-A/C-G loci to prevent strand ambiguities. SNPs with no chromosomal position were removed and coordinates were updated from rsIDs to a custom *Chr_position[b37]* ID to ensure a match across datasets. The minimum allele frequency was set to 1% (***--maf 0*.*01***) and missing genotypes were trimmed per individual and per locus to a maximum 5% (***--geno 0*.*05 --mind 0*.*05***).

Close kin (second degree relatives, kinship coefficient >0.087) were removed from all groups using the ***-king-cutoff 0*.*088*** flag. Outliers were detected iteratively using smartpca from the software package Eigensoft (42). We based removal on the first five eigenvectors for five iterations with a sigma threshold of 6 as in (43).

The final dataset comprised 164 100 SNPs, 52 populations and 806 individuals, of which 23 were BaPhuthi (n=2 South Africa, n=21 Lesotho) (Table_S 1).

#### 2.3.2 Data clustering and population structure

We investigated clustering in the data using ADMIXTURE v1.3.0 (44) and Principal Component Analysis (PCA). These approaches allowed us to confirm the absence of batch/chip effects or other merging artefacts and to compare the observed population structure to that previously reported. As neither analyses accounts for correlation between SNPs, we trimmed SNPs in linkage disequilibrium (45). We removed a locus from pairs with R2>0.7 for a 50bp frame with a 5bp sliding window (***PLINK --indep-pairwise 50 5 0*.*7*)** as per (20). The SNP count was reduced to 117 358 in these analyses.

ADMIXTURE was run for K values between 2 - 16 on the entire dataset, where K is the number of tested clusters. We used 10 replicates for each K and a 5-fold cross-validation error (CV) estimation with 100 bootstraps for standard errors (***-B100 --cv INPUTFILE*.*bed {2***..***16}***). The lowest CV error determined the optimum K-values (44). To identify common modes across replicates, we processed the output with the CLUMPPAK server (46) using default settings (LargeKGreedy algorithm, 2,000 random permutations). Results were visualised using ggplot2 (47) in R v.3.5.1 (48). The PCA was performed with PLINK (***--pca***) both on the entire data set and with a focus on relevant data from populations south of the African Forest Belt (∼ 5.6 S Latitude). We tested for significant differences in ADMIXTURE components between the focal populations using a Kruskal-Wallis rank-sum test, with pairwise differences identified with two post-hoc tests: the Baumgartner-Weiß-Schindler and the more conservative Nemenyi test, both from R package PMCMRplus (49).

### 2.4 Identifying parent populations and admixture dates

As a formal test for admixture in the history of the focal populations, we estimated the F_3_ indices (50; 51) in the form of F_3_(X,Y; Target population), where X and Y are potential source populations. Focal and reference populations were all included as possible sources. Negative F_3_ values (Z-score < -3) are considered indicative of a discordant tree relationship and in support of admixture (51). Estimates were made using Admixtools v.5.1 (52). The Lake Chrissie San were excluded from this analysis as they were represented by three individuals only.

We estimated the timing of admixture events by fitting exponential decay curves of linkage disequilibrium against increasing distances between SNP pairs, as implemented in MALDER (16; 53). Due to sample size, the Lake Chrissie San were excluded. We used an inter-generation time of 28 years, in line with other work (11; 20; 54). Events were estimated from 1960 CE, the mean date of birth of the BaPhuthi participants. Standard errors were estimated by jack-knifing over chromosomes and a Z-score was estimated by dividing the mean by the standard error.

## 3 Results

### 3.1 Data Clustering and Population Structure

Global population structure was explored using PCA and ADMIXTURE analysis. The patterns observed in the global PCA correspond well with published results on global diversity (Text_S 2, Figure_S 1, Figure_S 2). When focusing on the PCA among southern African groups (Figure 1), the first 5 PCs accounted for ∼6.5% of the total variation (Figure_S 3). Principal component 1 (PC 1, explaining ∼3.5% of the variation) separated KhoeSan populations from other African individuals (Figure 1). KhoeSan groups with known Eurasian admixture (‡Khomani, Nama and Karretjie) separated from the remaining Africans along PC 2 (1.01% of the variation). In PC 3 a separation between southern and northern communities emerged, particularly among the Bantu-speaking communities (Figure 1). On PC 4, the more western Africans are separated from eastern Africans (Figure 1).

On PCs 1 and 4, all the focal populations, except for the Lake Chrissie San, cluster close to one another. The Lake Chrissie San are distinctly closer to KhoeSan groups, in particular the Taa and Khoe-Kwadi groups. On PC 2 and 3 the Chrissie San are shifted more toward the KhoeSan which have no notable recent Eurasian admixture (e.g. G|ui & G||ana, Hai||om) than the Southern African ‡Khomani, Nama and Karretjie, but they are also shifted toward the Southern African SBE groups, showing some genetic similarity. Most of the BaPhuthi are at the extreme of the distribution of the Bantu-speaking populations along PC 3 (Figure 1). While this might be caused by a SNP-chip artefact, we do not see the same outlying position on other PCs (Figure_S 3) nor do we see an ADMIXTURE component in the unsupervised analyses which is unique to the BaPhuthi_LE (results below). Based on this, we suggest that the outlying position on PC 3 may instead reflect a BaPhuthi-specific change related to affinities to SBE-speakers.

The ADMIXTURE analysis of the global dataset paralleled the variation captured by the PCA described above, in line with previously published work (e.g. (55; 56)) reflecting divisions between global regions (Figure 2, Figure_S 4). We discuss here the results for K=9 (Figure 2) which is the largest K value with cross-validation errors *en par* with the lowest cross-validation errors (Figure_S 5). To simplify discussing the ADMIXTURE components we focus on those present in the focal populations, and refer to them based on the populations in which the component is at the highest proportion on average.

**Figure 2:**
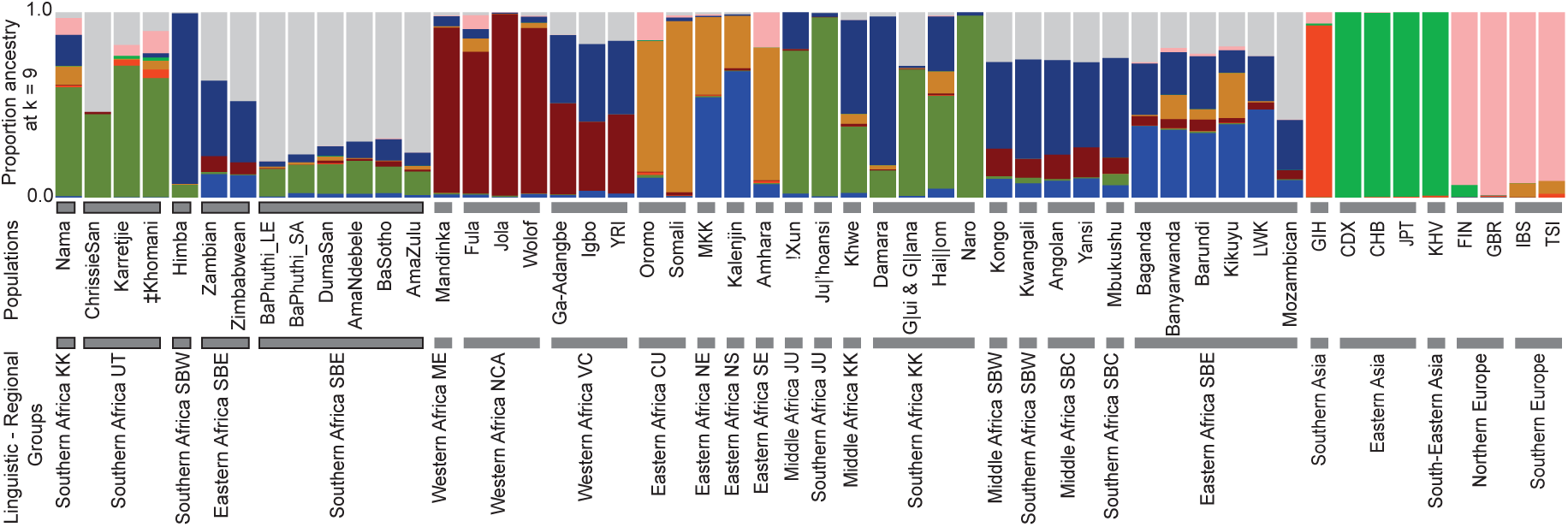
Population averaged ADMIXTURE proportions at K=9 represented as a stacked bar graph. Each colour represents a component. Samples are plotted in regional – linguistic groups. Population abbreviations are explained in Table S1.

The South African and Lesotho BaPhuthi had similar ADMIXTURE profiles. Based on this and their shared position on all PCA axes, we merged the data and from here forward we discuss the joint BaPhuthi data. Significant variation in ADMIXTURE components was found across our focal populations (BaPhuthi, Duma San, Lake Chrissie San, and the SBE) (Table_S 2, Kruskal-Wallis test KW=106.40, p<0.001). The disagreement between the post-hoc tests reflect different levels of conservativeness but where results were congruent, a pattern was observable.

The BaPhuthi profile was composed predominantly of two components but with minor contributions from an additional five components (Figure 2). The most predominant component was highest in the BaPhuthi (mean ∼81%, grey in Figure 2) but prevalent across the Southern African SBE and distinguished the Bantu language communities from other Africans. Among the focal groups, the Southern African SBE had significantly greater proportions when compared to the Southern African KhoeSan, the East African SBE and Southern African southern Bantoid, western Bantu language (SBW) communities (Table_S 2, Table_S 3, post-hoc tests).

The second largest component seen in the BaPhuthi (∼15%, dark green in Figure 2) was at its greatest in the Naro KhoeSan (98%), and occurred in other members of the focal group. In the BaPhuthi ADMIXTURE profile, the sum of the ‘Naro’ and ‘BaPhuthi’ components (95% ± 4%) was notably greater than the sum in the southern SBE (<90%) and Eastern SBE (<60%) (Table_S 3). Of the 22 Lesotho BaPhuthi, five (22%) had the ‘BaPhuthi’ and ‘Naro’ ADMIXTURE components make up >99% of their profile and, for 50% the two major components made up >95% of the profile. None of the SBE and only some KhoeSan groups had individuals with such profiles. The BaPhuthi clearly have a greater reduction in non-major components compared to other SBE. Beside the focal group, across the Southern African dataset only the Naro, Ju’|hoan, G|ui and G||ana had the sum of these two components comparable to the BaPhuthi (>95%). However in these populations the sum was largely driven by elevated ‘Naro’ proportions (the ratio ‘BaPhuthi’:’Naro’ was well below 1). In the BaPhuthi and the southern SBE the ratio was very similar (median ∼5.5 ± 1, Table_S 3) despite the variation in absolute values. The ratios were far greater than the Eastern African SBE where the ‘Naro’ component was virtually absent (1 ± 1%; mean ± s.d. Table_S 3).

The focal Eastern African SBE had significantly lower proportions of the ‘Naro’ component (< 1.5% Table_S 3) compare to the southern SBE (∼15 ± 2-3% Table_S 2, Table_S 3). The Zambians had greater proportions (1.4 ± 1.1% Table_S 2) than the Zimbabweans (0.6 ± 0.4% Table_S 3) despite being further North.

Of the minor components shared with the BaPhuthi, the component that is dominant in the Himba (‘Himba’ component, dark blue in Figure 2) showed the most variation within the focal Bantu-speakers. There was a lower proportion of the ‘Himba’ component in the Southern African SBE-speaking and !Ui – Taa speaking (UT) communities compared to the other Bantu-speaking groups from other regions and the Khoe-Kwadi speaking (KK) and Juu speaking (JU) KhoeSan (Table_S 3). Of the focal Southern African SBE, the BaPhuthi and AmaZulu had significantly lower proportions of the ‘Himba’ component compare to the BaSotho, Duma San and AmaNdebele, and much lower than the Zambians (41 ± 3%) and Zimbabweans (33 ± 3% Table_S 2). Admixture or drift likely differentiated the BaPhuthi and AmaZulu from the other southern SBE. The standard deviation of the population means for the ‘Himba’ component (0.29) was comparable to the standard deviation of the ‘BaPhuthi’ component (0.26), and both were greater than other components (<0.07, including the ‘Naro’). There is thus important variation of this western African component in the region. The co-occurrence in relatively similar amounts of the ‘BaPhuthi’, ‘Naro’, and ‘Himba’ components are suggestive of a shared history for the Southern African SBE populations.

The Nama are an exception within the region. They have distinctly elevated ‘Himba’ (17 ± 17% Figure 3) and ‘Somali’ (10 ± 5% Figure 3) components which are comparable to the other Khoe-Kwadi groups. Furthermore, the BaPhuthi, Duma San and Lake Chrissie San can be distinguished from the Nama, Karretjie and ‡Khomani by the lower levels of Eurasian components (light green and pink in Figure 2).

**Figure 3:**
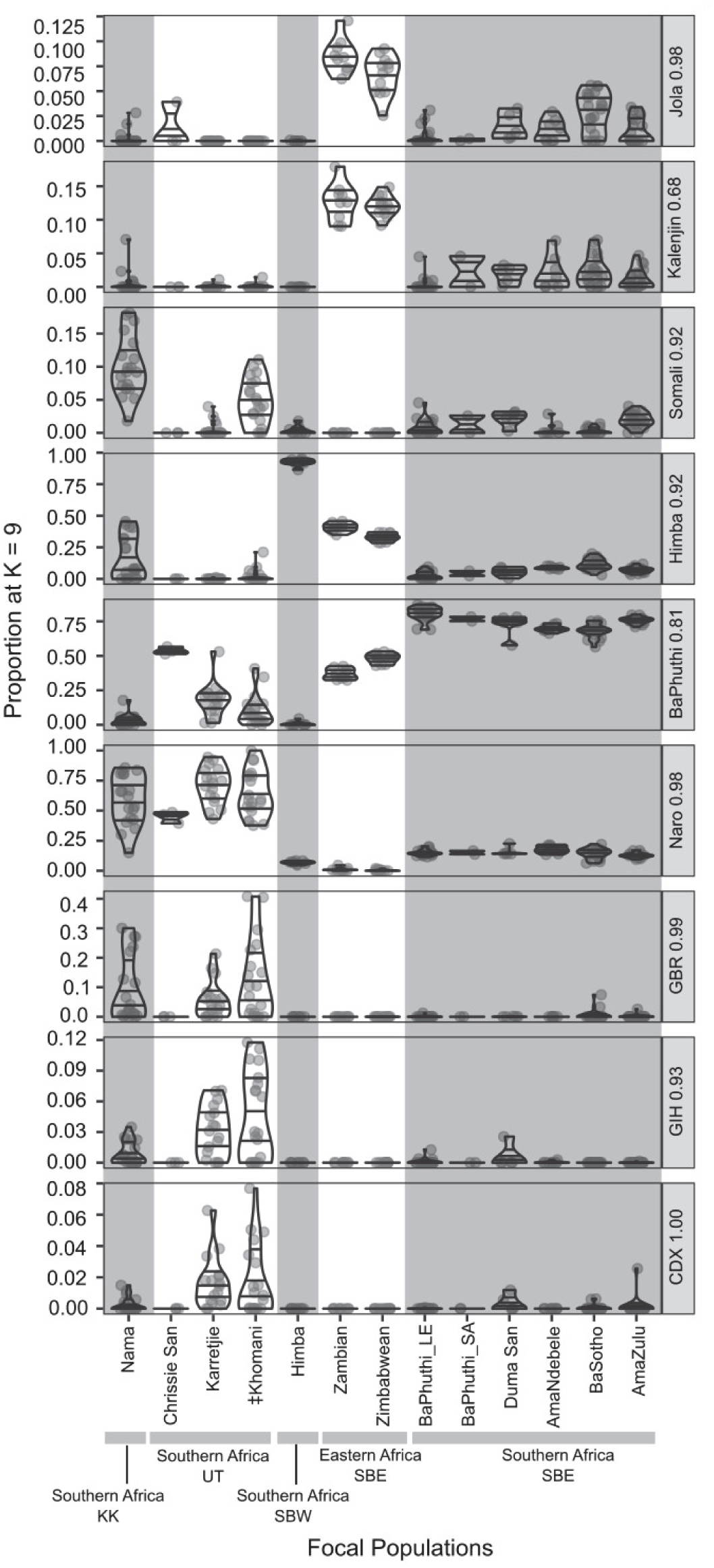
Variation in ADMIXTURE proportions for the focal populations at K=9. Dots represent individual data points and violin boxes show the range. Horizontal lines within the violins show median and inter-quartile ranges. Facet labels to the right of each row gives the population in which the component is at the highest proportion and the average value within that population. Note that the Y-axis range varies by facet.

### 3.2 Identifying parent populations and admixture dates

The overall similarity of the Southern African SBE was further supported by F_3_ admixture tests. In these populations, when the Ju’|hoansi paired with the west Africans (Yoruba or Igbo from Nigeria), the lowest Z-scores (Z< -10) were reported. In the Eastern African SBE, Z-score values recovered for the Zambians were below -8 and those of the Zimbabwean data were not statistically significant (Z > -3) (Table_S 4). Failure to detect the admixture in the Zimbabweans may reflect the lower proportion of KhoeSan ancestry as seen in the low ‘Naro’ ADMIXTURE component.

The influence of the East African pastoralist gene flow appears weaker in the Zambian, Zimbabwean and BaPhuthi as no admixture signals were detected for pairs involving non-Bantu East Africans and KhoeSan populations (Table_S 2). Signatures were detected in the Duma San, AmaZulu, AmaNdebele and BaSotho (Z-scores < -3) and for all the Southern African KhoeSan (Table_S 2).

Finally, only for Zambians and BaSotho there is evidence of an admixture event between two Bantu-speaking communities. A Central southern Bantu (SBC) or SBW admixture with a SBE group is detected in the Zambians. An Eastern Africa SBE – Southern African Nguni admixture is detected in the BaSotho.

As admixture across the SBE with the KhoeSan may not have be uniform, we tested to distinguish KhoeSan contributions which were distinct from the southern SBE using F_3_(KhoeSan,South African Bantu-speaking group,Target population). We obtained significant results for the BaSotho, Duma San and the AmaNdebele (Table_S 2) but not for the Zambian, Zimbabwean, AmaZulu and BaPhuthi. Thus the latter have less KhoeSan-related ancestry compared to the former. All the southern KhoeSan produced support for admixture with these pairs of sources, indicating that KhoeSan-related proportions and/or diversity were greater than the SBE.

We additionally tested for Eurasian ancestry not present in the KhoeSan by considering Eurasian - KhoeSan admixture. The Duma San, AmaNdebele and BaSotho again recovered a common significant F_3_ value, and in all cases the top results included GIH (Gujarati) or Eastern Asians. The BaPhuthi, AmaZulu, Zambian and Zimbabweans did not produce results to support such admixture. All the southern KhoeSan produced significant results for this test with the top scores for European – KhoeSan source pairs.

To provide chronological context to admixture events supported by the F_3_ results, we estimated admixture dates with linkage disequilibrium decay curves (MALDER, Table_S 5). Across the southern Bantu groups we detect admixture events over ∼100 generation ago. Here we focus on events younger than 12 Kya. Multiple admixture events were supported for the BaPhuthi, AmaZulu, Nama, Himba and Zambians. In the BaPhuthi we detected recent admixture between KhoeSan - European dating to ∼1786 CE. Such dates may correspond to colonial era European admixture as they are shared with events detected in the Karretjie, ‡Khomani and Nama (1808 – 1835 CE) which are known to be recent admixtures (6; 37; 57-59). Several of the southern SBE produced KhoeSan - European admixture dates distinctly older (1450 – 908 CE), indicating an event not tied to the 17^th^ century expansion of Europeans into Southern Africa. Several of the non-focal SBE populations produced similar dates (Table_S 5) and events detected in the Nama (252 CE – 139 BCE) were similar to the oldest of these dates. This suggests that the oldest Eurasian admixtures (∼908 BCE) may reflect pastoralist-related events.

Admixture between African populations were detected in the BaPhuthi (991 – 812 CE) and in the AmaZulu (1258 – 1143 CE) while older dates were detected for the Himba and Zambians (843 CE – 118 BCE), suggesting more recent admixture for the groups further south.

## 4 Discussion

The BaPhuthi have an oral history of descent from KhoeSan and a more recent narrative of the assimilation of refugees (29; 33). The results from our investigation show no support for a unique eastern KhoeSan ancestry to differentiate the BaPhuthi from other southern Bantu communities but the results do support a history distinct from other southern Bantu-speaking groups.

### 4.1 Close affinities to the southern Bantu communities

The BaPhuthi show strong genetic affinities to the communities from the surrounding region who speak southern Bantoid, eastern Bantu (SBE) languages. This was evident from the overlapping positions in the PCA, the similar ratio of ADMIXTURE components related to southern Bantu-speakers and KhoeSan (∼5:1) as previously reported (38; 60) and from the MALDER and F_3_ results. The profile was particularly similar to the AmaZulu in our analysis and in previous work (20; 34; 35). In contrast, there was a notable difference from the Duma San (SeSotho speaking) despite both sharing a narrative of recent KhoeSan descent (34).

While the BaPhuthi speak SePhuthi, a Southern/Lowland Ndebele language (40), we found the AmaNdebele shared more characteristics with the BaSotho and Duma San. This was despite that the AmaNdebele were geographically and linguistically nearer to the BaPhuthi and AmaZulu. Clearly the relationship among the Nguni is not simple. The divergence of the AmaNdebele from the other Nguni may result from non-uniform admixture between the Nguni and later arriving communities. The cultural differences of the Nguni and Sotho-Tswana provide some clues as the customs unique to the Nguni in Southern Africa have possible parallels in Rwanda. These include *hlonipha* (respectful etiquette) and a distinct dichotomy in social roles (e.g. between men tending to cattle while women tend to crops) (61), suggesting the arrival of an ‘intervening influence’ (possibly related to the BaSotho) between the regions.

The AmaNdebele showed support for additional Eurasian and KhoeSan admixture events not seen in the AmaZulu and BaPhuthi (Table_S 2, Table_S 3, Table_S 4) but in common with the BaSotho and Duma San. While the AmaNdebele migrated out of KwaZulu-Natal during the 1600s (61) and may have admixed during this migration, the East African/Eurasian ADMIXTURE components are too consistent across individuals to be from recent admixture. There was no support from F_3_ and MALDER (Table 1, Table_S 4). Instead the contribution may derive from admixture with another SBE group but here too there was no clear support.

**Table 1:**
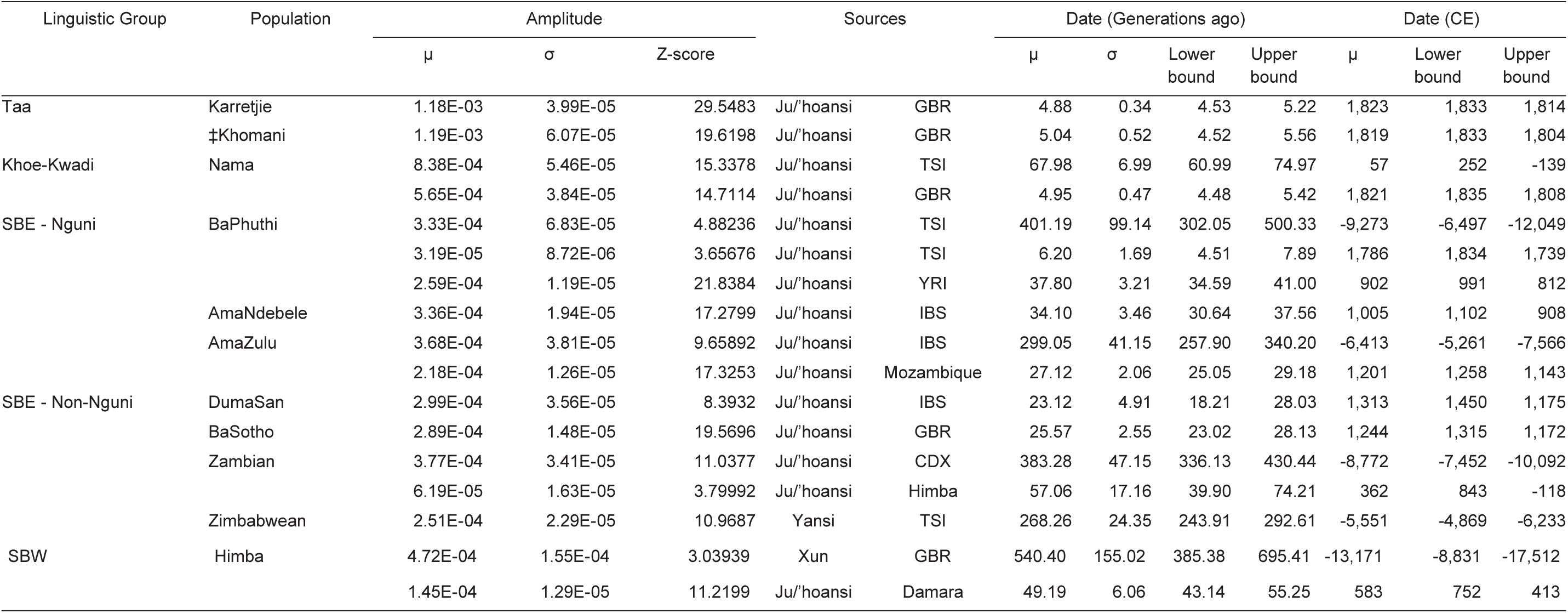
Admixture event dates estimated across populations based on linkage disequilibrium decay curves.

The Nguni are suspected to have been the earliest arrivals to Southern Africa of the present-day SBE (62) and their relatedness may give details to the regions early history. For example, the BaSotho may descend from a recent admixture between a southern SBE with a group from further North. Indeed the BaSotho could be modelled as F_3_(BaPhuthi,Zambians,BaSotho) or F_3_(Duma San,Zimbabweans,BaSotho) (but no other combination of these, Table_S 4). This result would relate to the elevated western Bantu contribution in the Zambians and Duma San (increased ‘Himba’ and ‘Naro’ ADMIXTURE components, Table_S 3). The detected admixture in the BaSotho and Duma San dating to 1450 – 1173 CE (Table 1) may reflect a late iron-age (LIA) contact. Both speak SeSotho, a language related to SeTswana, TshiVenda and Makua, and found predominantly in Mozambique and Botswana (40; 63; 63). With our result, this could suggests a shared history further North and that the ‘proto-BaSotho’ migrated southward.

The origin of the SBW component in the BaSotho and Duma San may relate to that in the Zambians. This contribution may be tied to the establishment of Naviundu pottery tradition in southern Congo and ultimately a possible western Bantu source (64). Our MALDER dates for such an event, 843 CE – 118 BCE, coincide with the establishment of early iron-age (EIA) in the region (Madingo-Kayes, ∼2 CE (65)).

### 4.2 A Complex KhoeSan Descent

In common with other Bantu-speaking communities, the BaPhuthi retain oral history of KhoeSan descent and culture which is reflective of KhoeSan influence (8; 34; 66). We found support for a common origin of the KhoeSan heritage with that of the southern SBE and southern KhoeSan but no support for a unique KhoeSan descent in the BaPhuthi. In some SBE (e.g. Zambians), KhoeSan descent was detected which was distinct from the southern SBE. The BaPhuthi were not among these groups, contrary to what was anticipated from the ‘essentialistic’ understanding of ‘Bushman’ descent. Moreover, the component prevalent in the BaPhuthi was most prevalent in the Karretjie, ‡Khomani and Chrissie San suggesting a regional affinity.

The KhoeSan ADMIXTURE component, referred to as the ‘Naro’ component, is seen in the BaPhuthi and the southern SBE, Duma San and Lake Chrissie San. The southern SBE lacked a clear difference from the southern KhoeSan based on the F_3_ statistics and among the southern SBE, the ‘BaPhuthi’:’Naro’ ADMIXTURE ratios were remarkably consistent. This is particularly noteworthy considering the variation in linguistic affinities and other ancestral components of the southern SBE. These results supports the proposed common source and event for the KhoeSan admixture in the region (11; 39). When compared to other SBE, the southern SBE have elevated KhoeSan components. This is often argued as the result of demographic dynamics when the EIA Bantu language expansion progressed across in southern Africa (39; 67; 68). The change in environmental conditions would have slowed the rate of population growth and allowed for greater admixture with local populations (69) leading to greater KhoeSan ancestry.

If we assume that the KhoeSan affinities of the BaPhuthi indeed reflect a recently acquired ancestry from the Maloti-Drakensberg, these ancestors would then have had shared affinities with other groups across the region. The broad relatedness across the region is possible as the ancestral languages of the groups in which the ‘Naro’ ADMIXTURE component was largest (G|ui and G||ana, the Karretjie (likely |Xam language), the ||Xegwi (Chrissie San ancestors)) belonged to the !Ui branch of the Tuu family (34). The languages spoken around the Drakensberg are likely to have also been Tuu (70).

The southern SBE and southern KhoeSan appear to have a common ancestral KhoeSan population but the admixture event is not likely to be shared. The amount of KhoeSan heritage and the ratios of ‘BaPhuthi’:’Naro’ were notably different between the groups. The consistency of the ratios in the southern SBE would be explained most parsimoniously by a common admixture event and then dispersal across the region while the variation among the southern KhoeSan suggests independent admixture events. This would then suggest a common ancestral ‘proto-southern SBE’ who drifted or admixed to give rise to the present-day SBE diversity.

Some traits make the BaPhuthi data peculiar from other SBE. Of particular interest is that the Bantu-related ‘BaPhuthi’ and ‘Naro’ ADMIXTURE components are at higher proportions compared to other southern SBE, yet several Bantu-related components are frequently absent from BaPhuhti (Figure_S 4). The ‘BaPhuthi’ and ‘Naro’ components sum to >95% for almost half of the BaPhuthi but none of the other southern SBE had any individuals with such high sums (median total of the two components ∼85 – 89%, Figure_S 4). Furthermore, components such as the ‘Himba’, ‘Kalenjin’ and ‘Jola’ which were almost ubiquitous in the southern SBE, were frequently absent form the BaPhuthi. The BaPhuthi have apparently diverged even from their closet genetic kin, the AmaZulu. For perspective, consider that only the Lake Chrissie San, ǂKhomani, Karretjie, and G|ui and G||ana had individuals with similar profiles (i.e. the sum of the two components >90%).

An independent loss of the same minor components in the BaPhuthi, southern KhoeSan and G|ui and G||ana seems unlikely considering the large differences in the proportions of the two major components. This is particularly for the BaPhuthi where >95% of the ancestry in some individuals is southern SBE related. However, the outlying position of the BaPhuthi on PC3 (Figure_S 1) may reflect a bottleneck related to southern Bantu-speaking and KhoeSan affinities. The result together are ambiguous and require further work.

A separate (re)introduction of the minor components is supported in several SBE groups. The AmaZulu and BaPhuthi could not be modelled as an admixture of East Africans and a KhoeSan group with F_3_, suggesting that the East African ancestry detected in the BaSotho, Duma San and AmaNdebele (e.g. the Asian admixture on F_3_, ‘Somali’, ‘Kalenjin’, ‘Himba’ ADMIXTURE components, Figure 2, Table_S 4) may reflect subsequent admixture rather than recent drift in the Nguni. The Duma San acknowledged recent Bantu-speaker ancestors (34) which may be the source of the (re)introduced components. The BaPhuthi too acknowledge recent SBE admixture, and it is evident in some individuals, but it is clearly not ubiquitous in the region and may not have left a detectable signal.

The possible re-introduction parallels the argument raised by (64) in that there is a pattern for an early arriving iron age Bantu-related community, possibly Kalundu Tradition facies (here the ‘Proto-southern SBE’ ADMIXTURE profile of the BaPhuthi) and then a replacement or assimilation by LIA groups possibly derived from EIA Urewe Tradition facies (marked by the addition of the ‘Himba’, ‘Kalenjin’ and ‘Jola’ ADMIXTURE components as in Banyarwanda, Barundi). MALDER date estimates for admixture between KhoeSan and non-KhoeSan Africans (991 CE – 118 BCE in the Zambians, Himba & BaPhuthi) pre-date the arrival of the LIA Bantu expansion into south-eastern Africa (18; 21). While dates detected in the other southern SBE (1102 – 908 CE AmaNdebele; 1450 – 1143 CE AmaZulu, Duma San, BaSotho) for admixture between KhoeSan and non-KhoeSan could be related to the second millennium CE (71).

Our results can thus support an arrival of Moloko pottery/Sotho-Tswana from further north (admixture detected in the BaSotho and Duma San) while the impact of the arrival of the Blackburn pottery style in KwaZulu Natal is unclear without regional contrasts.

We found no ‘ancient’ distinction of the BaPhuthi from the SBE in KhoeSan ancestry, but the BaPhuthi may have recently incorporated AmaTola (32; 33). The AmaTola reportedly incorporated Khoikhoi during the Frontier wars (1779 – 1878 CE) (8) and in the AmaTola descendants, the Duma San, we do see greater proportions of ADMIXTURE components in common with the Nama (a Khoikhoi group). While Khoikhoi pastoralists are recorded along the western parts of South Africa (15) and left cognates in the Nguni languages (62), the extent of their range eastward is unclear (72). The Nama have genetic ancestry indicating an admixture between a southern KhoeSan group and a Eurasian group related to the arrival of pastoralism in the region (11; 59) possibly 252 CE – 139 BCE (Table 1). If we assume that the East African components identified in the southern SBE entered during the migration through East Africa into Southern Africa (71; 73), then the absence of elevated East African ancestry in the BaPhuthi would indicate that the assimilated ‘Khoikhoi’ were perhaps culturally pastoralists but not of Khoe-Kwadi descents or that the component has since been lost to genetic drift.

### 4.3 The Heterodoxy

Turning our attention to more recent admixture, we did not find support for Eurasian contributions for the BaPhuthi. The reported history of the BaPhuthi often refers to the amalgamation of communities, amongst them were the predecessors to the Duma San, the AmaTola, and possibly recent slave descendants (29). The F_3_ results show no support for Eurasian admixture which cannot be explained with ancestry already in a KhoeSan group (e.g. the Ju’|hoansi). The same result is found for the AmaZulu, Zambians and Zimbabweans, much unlike the young admixtures supported by MALDER and F_3_ in the Karretjie, ǂKhomani and Nama (1834 – 1804 CE). While an admixture in the BaPhuthi was detected by MALDER (Ju’|hoansi – TSI, 1833 – 1739 CE), the absence of ADMIXTURE components and support from F_3_ means this event may reflect admixture with a pastoralist-related group such as the Khoikhoi.

Considering that the AmaTola and BaPhuthi were at their names-sake height during the mid 19th century (200 years ago) (8), the introduction of Eurasian ancestry should be detected in a sample of 23 individuals. While this doesn’t exclude historic assimilation of Eurasians into the broader BaPhuthi community, it does indicate that it was not a wide-spread phenomenon. None of our participants disclosed non-African descent such that our expectation of a recent Eurasian contribution stems entirely from literature (29; 32) and does not reflect local history.

In this work we investigated to what extent the oral history of BaPhuthi as KhoeSan descendants is reflected in their genetics. The genetic composition of BaPhuthi harbour signals for an interesting connection to the early arrival of the Bantu languages but we could not support a unique eastern KhoeSan contribution. In the case of the BaPhuthi and the previously investigated Duma San, we find some support for KhoeSan descent but not necessarily in the ‘essentialistic’ understanding of an eastern KhoeSan. Such essentialistic reading in historic literature can create misconstrued narratives of ethnic/biological distinctions. The high status attained by KhoeSan Shaman and the pride taken in the KhoeSan means of semi-nomadic subsistence (8) has entrenched the importance of KhoeSan heritage in the collective memory of the BaPhuthi and perhaps without necessarily reflecting recent assimilation of some remnant ‘essentialistic’ KhoeSan group. The colonial era references to ‘San’ and ‘Bushman’ may reflect ambiguous discussions of non-sedentary polities and practices of raiding and mixed subsistence. Equally challenged here are the accounts of assimilation of Eurasian runaway slaves and servants, and people of mixed-race into the developing BaPhuthi as we see no evidence for widespread recent Eurasian admixture.

## 5 Appendices

## 6 Supplementary Data

Supplemental Data includes five figures, six tables, and Supplemental Material and Methods can be found with this article online at [xxxxx].

## 7 Declaration of Interests

The authors declare no competing interests.

8 Acknowledgement

We thank the all the participants for their time and engagement. R.J.D thanks the South African National Research Foundation, the Commonwealth Commission in the U.K., Santander Travel grant and the Oppenheimer Memorial Trust for support. The project was funded by a Commonwealth Commission in the U.K. and Boise Trust Fund award to R.J.D. The funders had no role in study design, data collection, analyses, decision to publish, preparation of the manuscript or the outcome. We thank Alessandro Raveane for useful discussions and assistance. All data analyses were performed at the ILIFU High-Performance Computer Centre of the University of the Western Cape, South Africa (https://www.ilifu.ac.za/).

## 9 Web Resources

The 1000 Genomes Project: http://www.internationalgenome.org/home

Human Evolutionary Genetics Group Data: https://capelligroup.wordpress.com/data/

Schlebusch Data: http://jakobssonlab.iob.uu.se/data

African Genome Variation Project: https://ega-archive.org/dacs/EGAC00001000237

Pickrell Data: https://reich.hms.harvard.edu/datasets

## 10 Data and Code Availability

The datasets generated during this study are available at [name of repository] [accession code/web link]. There are restrictions to the availability of the dataset due to participant confidentiality agreements. To access the data please contact the corresponding authors.

