## Supplemental Information for "Genetic heritage of the BaPhuthi highlights an over ethnicised notion of ‘Bushman’ in the Maloti-Drakensberg, Southern Africa"

#### Table of Figures

Figure_S 1: Principal component analysis of all data. Focal populations are indicated by black overlain symbols. Colours indicate regional and linguistic divisions. Linguistic abbreviations: Mande - ME, North-Central Atlantic - NCA, Volta-Congo - VC, Cushtic – CU, Nilotic Eastern - NE, Nilotic Southern- NS, Semitic - SE, Southern Bantu Western – SBW, Southern Bantu Central Western – SBC, Southern Bantu Eastern – SBE, Juu KhoeSan – JU, Khoe-Kwadi KhoeSan – KK, Ui! and Taa KhoeSan – UT........5
Figure_S 2: Principal component analysis of the global dataset, plotting only the South African data. Focal populations are indicated by black overlain symbols. Colours indicate regional and linguistic divisions. Linguistic abbreviations: Mande - ME, North-Central Atlantic - NCA, Volta-Congo - VC, Cushtic – CU, Nilotic Eastern - NE, Nilotic Southern- NS, Semitic - SE, Southern Bantu Western – SBW, Southern Bantu Central Western – SBC, Southern Bantu Eastern – SBE, Juu KhoeSan – JU, Khoe-Kwadi KhoeSan – KK, Ui! and Taa KhoeSan – UT........7
Figure_S 3: Principal component analysis of Southern African data. Focal populations are indicated by black overlain symbols. Colours indicate regional and linguistic divisions. Linguistic abbreviations: Mande - ME, North-Central Atlantic - NCA, Volta-Congo - VC, Cushtic – CU, Nilotic Eastern - NE, Nilotic Southern- NS, Semitic - SE, Southern Bantu Western – SBW, Southern Bantu Central Western – SBC, Southern Bantu Eastern – SBE, Juu KhoeSan – JU, Khoe-Kwadi KhoeSan – KK, Ui! and Taa KhoeSan – UT........9

### 1 Supplemental Text

*Text_S 1: Discussion on the use of population 'labels'*

On the backdrop of colonial era racism, slavery, and genocide, people are rightfully sensitive about the words used to discuss them (1-3). Participants in research should feel respected and that their sentiments are understood both to encourage ongoing public trust and for growth within the scientific community.

Terms used in our paper, such as ‘Hottentot’, ‘Khoisan’, ‘Bantu’ and ‘Bushman’ have a negative connotation to many who have been presently or historically referred to as such (4; 5) . All identifiers or ethnic labels carry some negative connotation because conflict with external communities is unavoidable. As such it is very challenging to find non-offensive terms when referring to communities and moreover when attempting to co-ordinate the use of terms across regions where histories are different. For example, while is South Africa the term ‘Bushman’ is strongly derogatory, some Kalahari hunter-gatherers prefer the name ‘Bossiesmans’ (Afrikaans for ‘Bushmen’) (5).

Despite ongoing debate on the use of ethnic labels, some recommendations are available (6; 7). The Working Group of Indigenous Minorities in Southern Africa and the South African San Institute now represent the Indigenous communities in the region. Following the 2003 African Human Genome Initiative conference the two institutions declared a preference for their individual community names or collectively as San. The term Khoe–San was optional when discussing the San and Khoikhoi as a collective. ‘Khoisan’ was first coined by anthropologist Leonard Schultze to refer to the pastoralist Khoi and the hunter-gatherer San. As the term persists in linguistics for the broad and diverse language area, we chose to use it in our work (modified as ‘KhoeSan’). We do not assume that those included are linguistically, culturally or genetically homogenous.

The term Bantu means ‘people’ in the Bantu languages. While the term is not offensive across Africa, in Southern Africa the term can be seen as offensive because of its use by the Apartheid regime in South Africa. As such we opted for using the term solely in the linguistic context of referring to the collection of people who speak the Bantu languages, i.e. Bantu speaking communities.

Lastly, as our paper attempts to investigate possible genetic distinctions between ‘vanished’ communities who are know mostly, if not entirely, from historic texts, we necessarily need to consider the terminology used in historic texts. Many of these terms have been identified by communities as derogatory and should be avoided (6; 7). As we do not want to make the field more complicated by introducing new terms, we chose to use the term ‘Bushman’ but solely to make it clear to which historic references we are discussing.

We are not referring to any contemporary people, participants or communities as ‘Bushman’ nor do we consider the term ‘Bushman’ to be a legitimate way to refer to people except where it may be the preferred term (e.g. in the Kalahari).
Furthermore, our results show that the notion of a genetically distinct ‘Bushman’ community

is unsupported and the term may have no value beyond discussing the use of the word in colonial literature.

*Text_S 2: Discussion of global PCA results*

The first 5 PCs accounted for ~11% of the total variation. Principal component 1 (PC1, explaining ~6% of the variation) separated African from Eurasian individuals (Figure_S 1, Figure_S 2). Principal component 2 (~2% of the variation) separated Eastern and Western Eurasians (Figure_S 1, Figure_S 2). Along PC3 (2% of the variation) the KhoeSan individuals are separated from the non-KhoeSan Africans. Principal components 1 and 3 show the BaPhuthi from Lesotho and South Africa plot close to the southern Bantu-speaking populations (e.g. AmaZulu) and the Duma San. The Lake Chrissie San are distinctly closer to the Naro, Ju/'hoansi and G|ui and G||ana compared to Duma San and BaPhuthi individuals. The recently admixed Southern KhoeSan groups (‡Khomani, Nama and Karretjie) are spread toward Eurasian groups along PC1 which is not seen for the BaPhuthi nor Lake Chrissie or Duma San.

An East African component found in the horn of Africa (Somali, Oromo, Amhara) is identified by PC4 (Figure_S 1). On this PC the BaPhuthi, Duma San and southern Bantu-speaking groups are shifted toward the KhoeSan groups, away from East Africans. On PC5, West Africans are separated from East and Southern Africans and here we see the BaPhuthi and Duma San are at the extreme end of the Southern African cline, beyond the other southern Bantu-speakers (Figure_S 1, Figure_S 2). The Lake Chrissie San are closer to the Hai||om than the BaPhuthi and Duma San on PC5.

While accounting for <1% of the variation, PC 7 and 9 highlights two interesting affinities in the BaPhuthi. On PC7 the BaPhuthi plot at the extreme end near the Southern African KhoeSan, as opposed to the Juu and Khoe-Kwadi at the other end. The Southern African SBE are also shifted in this direction but the BaPhuthi are well beyond other South African KhoeSan descendants (Duma San, Lake Chrissie San).
The PC9 distinguishes a Western/Juu (e.g. Xun, Ju/'hoansi) KhoeSan component from a Southern/Taa (‡Khomani, Karretjie) and Khoe-Kwadi component (Nama etc.) (Figure_S 1, Figure_S 2). While the Southern African SBE (including the Duma San) are shifted toward the Western/Juu compared to the Eastern African SBE, the BaPhuthi from Lesotho are shifted further. The Lake Chrissie San, in contrast, are slightly off centre toward the Southern/Taa groups.

The outlying position of the BaPhuthi_LE along PC5 and 7 is unlikely due to a SNP-chip artefact as we do not see a unique BaPhuthi_LE component in the unsupervised ADMIXTURE analyses (Figure_S 4). This suggests that the outlying position instead reflects possible another evolutionary cause within the BaPhuthi related specifically to a reduction in diversity in the KhoeSan (PC9) and southern Bantu-speaker affinities (PC5).

### 2 Supplemental Figures

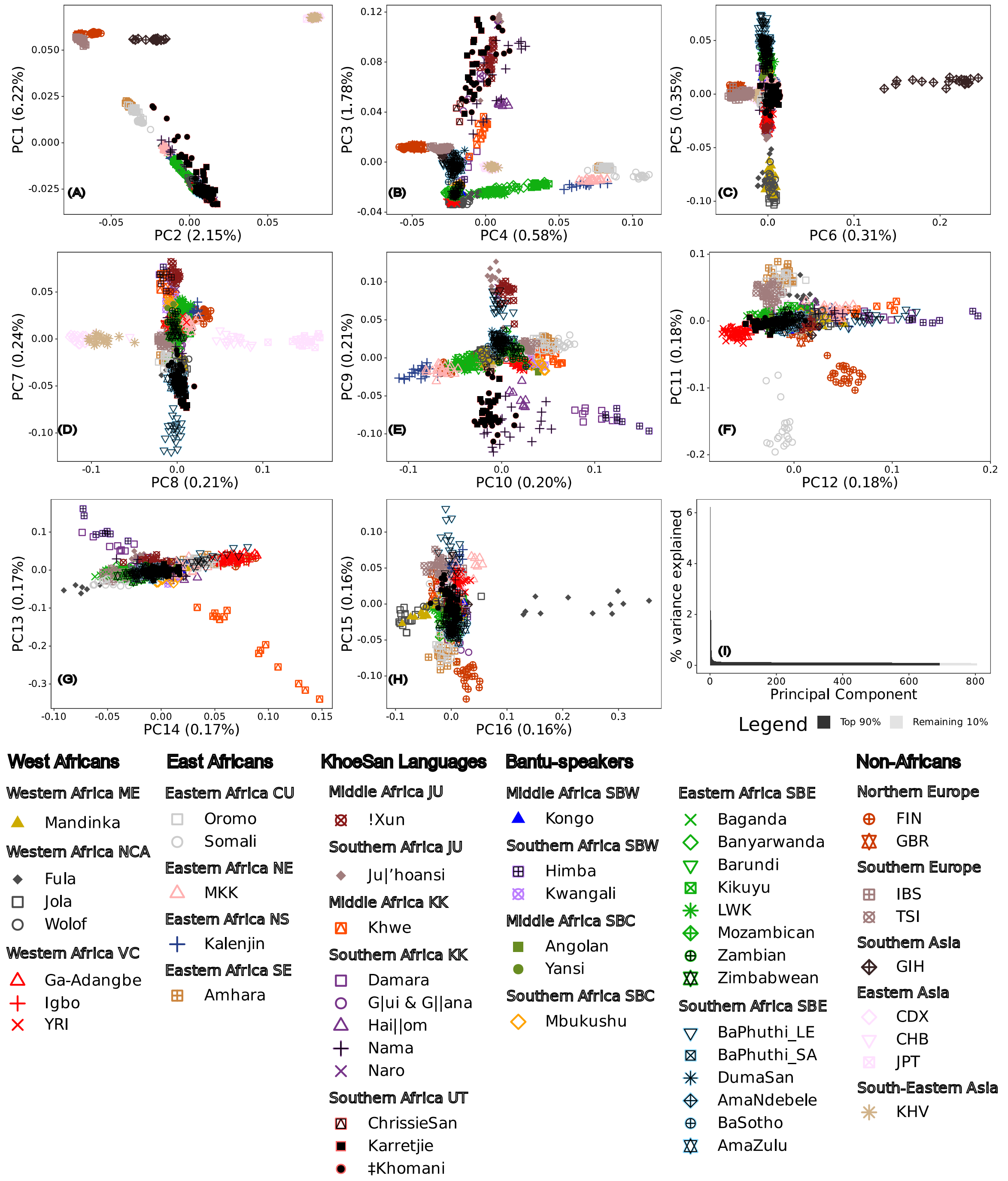

*Figure_S 1: Principal component analysis of all data used. Focal populations are indicated by black overlain symbols. Colours indicate regional and linguistic divisions. Linguistic abbreviations: Mande - ME, North-Central Atlantic - NCA, Volta-Congo - VC, Cushtic – CU, Nilotic eastern - NE, Nilotic southern- NS, Semitic - SE, southern Bantoid western Bantu– SBW, southern Bantoid central western Bantu– SBC, southern Bantoid Eastern Bantu– SBE, Juu KhoeSan – JU, Khoe-Kwadi KhoeSan – KK, Ui! and Taa KhoeSan – UT.*

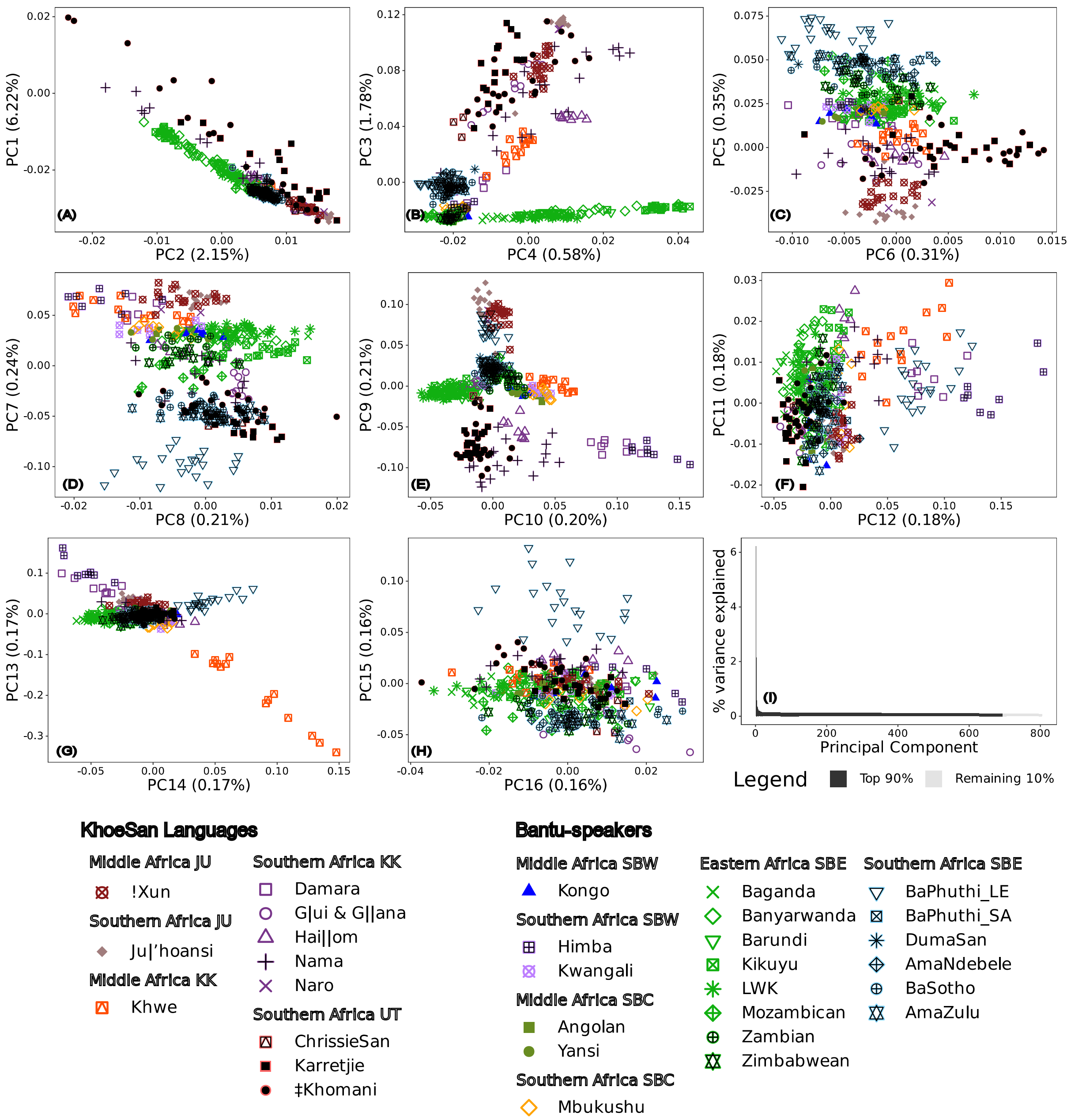

*Figure_S 2: Principal component analysis of all data used, plotting only the South African samples. Focal populations are indicated by black overlain symbols. Colours indicate regional and linguistic divisions. Linguistic abbreviations: Mande - ME, North-Central Atlantic - NCA, Volta-Congo - VC, Cushtic – CU, Nilotic eastern - NE, Nilotic southern- NS, Semitic - SE, southern Bantoid western Bantu– SBW, southern Bantoid central western Bantu– SBC, southern Bantoid Eastern Bantu– SBE, Juu KhoeSan – JU, Khoe-Kwadi KhoeSan – KK, Ui! and Taa KhoeSan – UT.*

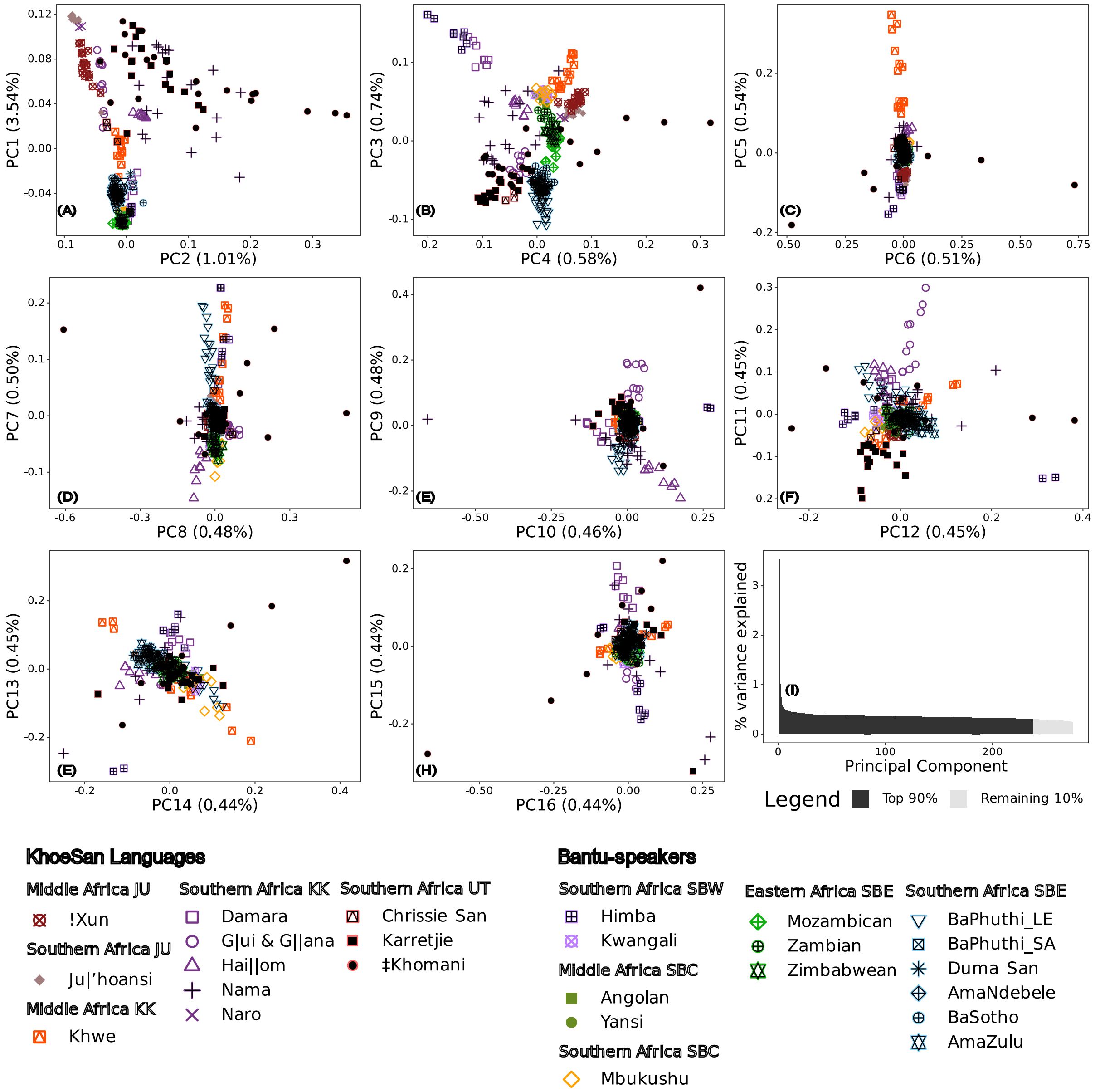

*Figure_S 3: Principal component analysis of Southern African data. Focal populations are indicated by black overlain symbols. Colours indicate regional and linguistic divisions. Linguistic abbreviations: Mande - ME, North-Central Atlantic - NCA, Volta-Congo - VC, Cushtic – CU, Nilotic eastern - NE, Nilotic southern- NS, Semitic - SE, southern Bantoid western Bantu– SBW, southern Bantoid central western Bantu– SBC, southern Bantoid Eastern Bantu– SBE, Juu KhoeSan – JU, Khoe-Kwadi KhoeSan – KK, Ui! and Taa KhoeSan – UT.*

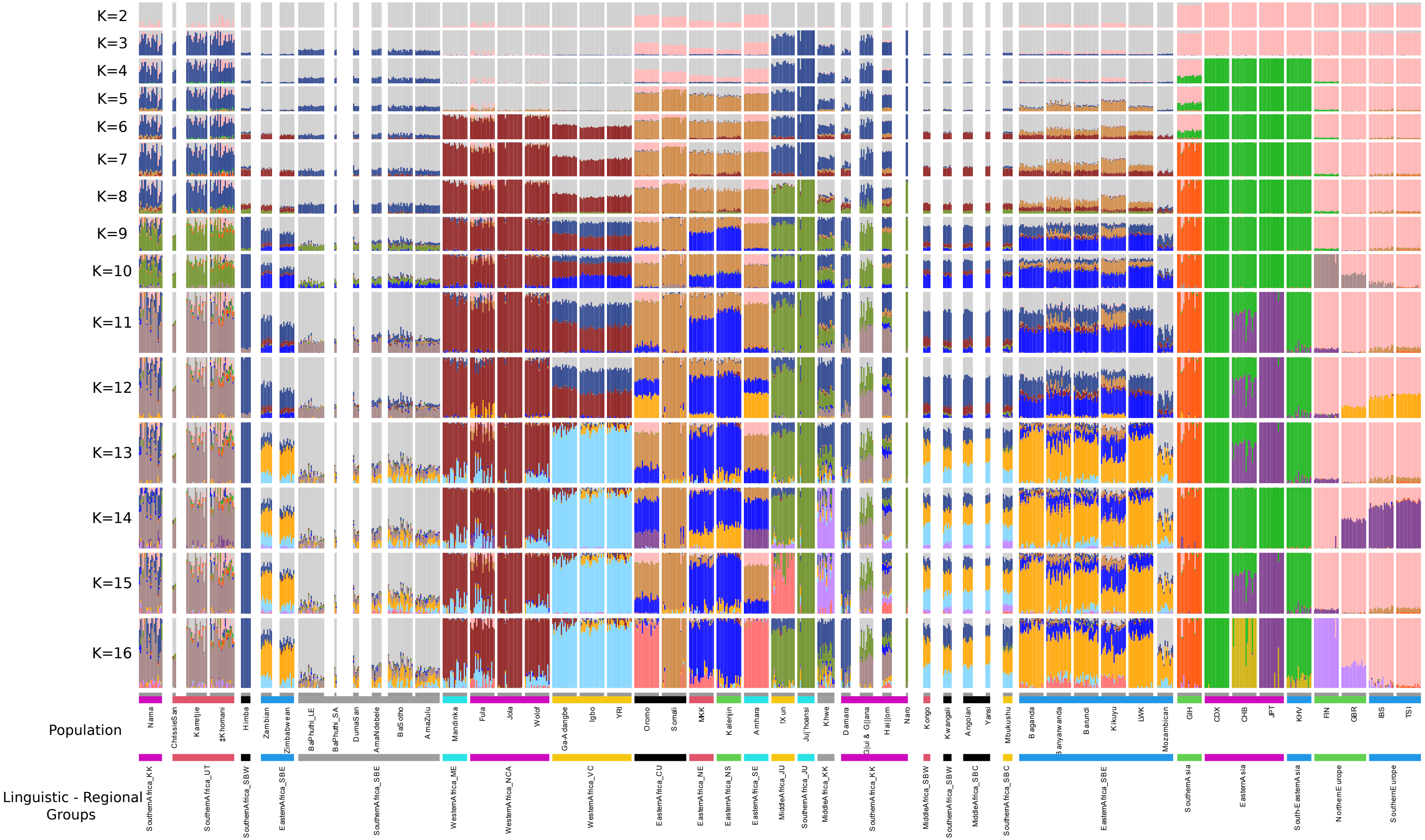

*Figure_S 4: ADMIXTURE proportions for K=2 ...16 represented as a stacked bar graph for each individual. Each colour represents a component. Samples have been grouped by Linguistic-geographic regions. Population abbreviations are explained in Table_S 1.*

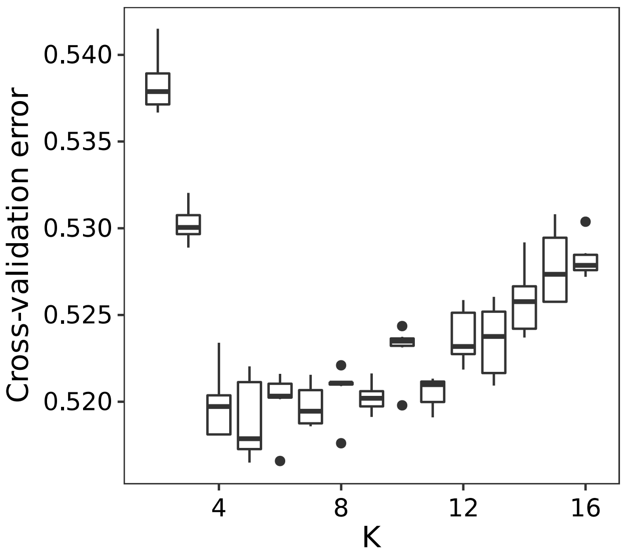

*Figure_S 5: Changes in cross-validation error for K=2..16 with 10 repeated runs each. Dots indicate outlying values. Median value indicated by the line within each box, and the inter-quartile range is indicated by the extent of the boxes.*

#### 2.1 Supplemental Tables

*Table_S 1: List of data included in this paper. Sample sizes after quality control steps (n) are indicated. Language information taken from Glottolog v4.*

| Reference or focal | Group | Reference | Continent | Region | Group Label | Language Family | Language Subfamily | Language Branch | n |
|---|---|---|---|---|---|---|---|---|---|
| Focal | Zambian | (8) | Africa | Eastern Africa | EasternAfrica_SBE | Atlantic-Congo Family | Volta-Congo | Benue-Congo | 9 |
| | Zimbabwean | (8) | Africa | Eastern Africa | EasternAfrica_SBE | Atlantic-Congo Family | Volta-Congo | Benue-Congo | 12 |
| | Nama | (9) | Africa | Southern Africa | SouthernAfrica_KK | Central/Khoe | Khoekhoe | Northern | 19 |
| | AmaZulu | (10) | Africa | Southern Africa | SouthernAfrica_SBE | Atlantic-Congo Family | Volta-Congo | Benue-Congo | 20 |
| | BaSotho | (10) | Africa | Southern Africa | SouthernAfrica_SBE | Atlantic-Congo Family | Volta-Congo | Benue-Congo | 20 |
| | AmaNdebele | (8) | Africa | Southern Africa | SouthernAfrica_SBE | Atlantic-Congo Family | Volta-Congo | Benue-Congo | 8 |
| | Duma San | (11) | Africa | Southern Africa | SouthernAfrica_SBE | Atlantic-Congo Family | Volta-Congo | Benue-Congo | 5 |
| | BaPhuthi_LE | This paper | Africa | Southern Africa | SouthernAfrica_SBE | Atlantic-Congo Family | Volta-Congo | Benue-Congo | 21 |
| | BaPhuthi_SA | This paper | Africa | Southern Africa | SouthernAfrica_SBE | Atlantic-Congo Family | Volta-Congo | Benue-Congo | 2 |
| | Himba | (12) | Africa | Southern Africa | SouthernAfrica_SBW | Atlantic-Congo Family | Volta-Congo | Benue-Congo | 8 |
| | ‡Khomani | (9) | Africa | Southern Africa | SouthernAfrica_UT | Southern/Tuu/Taa-!Ui | !Ui and Taa | Taa | 20 |
| | Karretjie | (9) | Africa | Southern Africa | SouthernAfrica_UT | Southern/Tuu/Taa-!Ui | !Ui and Taa | !Ui | 17 |
| | Chrissie San | (11) | Africa | Southern Africa | SouthernAfrica_UT | Southern/Tuu/Taa-!Ui | !Ui and Taa | !Ui | 3 |
| Global Reference | Oromo | (10) | Africa | Eastern Africa | EasternAfrica_CU | Afro-Asiatic Family | Cushitic | East Cushtic | 20 |
| | Somali | (10) | Africa | Eastern Africa | EasternAfrica_CU | Afro-Asiatic Family | Cushitic | East Cushtic | 20 |
| | MKK | (13) | Africa | Eastern Africa | EasternAfrica_NE | Nilotic Family | Eastern Nilotic | Teso-Lotuxo-Maa | 20 |
| | Kalenjin | (10) | Africa | Eastern Africa | EasternAfrica_NS | Nilotic family | Southern Nilotic | Kalenjin | 20 |
| | LWK | (13) | Africa | Eastern Africa | EasternAfrica_SBE | Atlantic-Congo Family | Volta-Congo | Benue-Congo | 20 |
| | Baganda | (10) | Africa | Eastern Africa | EasternAfrica_SBE | Atlantic-Congo Family | Volta-Congo | Benue-Congo | 20 |
| | Banyarwanda | (10) | Africa | Eastern Africa | EasternAfrica_SBE | Atlantic-Congo Family | Volta-Congo | Benue-Congo | 20 |
| | Barundi | (10) | Africa | Eastern Africa | EasternAfrica_SBE | Atlantic-Congo Family | Volta-Congo | Benue-Congo | 20 |
| | Kikuyu | (10) | Africa | Eastern Africa | EasternAfrica_SBE | Atlantic-Congo Family | Volta-Congo | Benue-Congo | 20 |
| | Mozambican | (8) | Africa | Eastern Africa | EasternAfrica_SBE | Atlantic-Congo Family | Volta-Congo | Benue-Congo | 13 |

| | | | | | | | | |
|---|---|---|---|---|---|---|---|---|
| Amhara | (10) | Africa | Eastern Africa | EasternAfrica_SE | Afro-Asiatic Family | Semitic | West Semitic | 20 |
| !Xun | (9) | Africa | Middle Africa | MiddleAfrica_JU | Northern/!Kung | Ju | | 19 |
| Khwe | (9) | Africa | Middle Africa | MiddleAfrica_KK | Central/Khoe | Kalahari Khoe | Western | 14 |
| Angolan | (8) | Africa | Middle Africa | MiddleAfrica_SBC | Atlantic-Congo Family | Volta-Congo | Benue-Congo | 8 |
| Yansi | (8) | Africa | Middle Africa | MiddleAfrica_SBC | Atlantic-Congo Family | Volta-Congo | Benue-Congo | 4 |
| Kongo | (8) | Africa | Middle Africa | MiddleAfrica_SBW | Atlantic-Congo Family | Volta-Congo | Benue-Congo | 6 |
| Ju\|'hoansi | (9) | Africa | Southern Africa | SouthernAfrica_JU | Northern/!Kung | Ju | | 14 |
| Damara | (14) | Africa | Southern Africa | SouthernAfrica_KK | Central/Khoe | Khoekhoe | Northern | 8 |
| Hai\|\|om | (14) | Africa | Southern Africa | SouthernAfrica_KK | Central/Khoe | Khoekhoe | Northern | 8 |
| Naro | (12) | Africa | Southern Africa | SouthernAfrica_KK | Central/Khoe | Kalahari Khoe | Western | 2 |
| G\|ui & G\|\|ana | (9) | Africa | Southern Africa | SouthernAfrica_KK | Central/Khoe | Kalahari Khoe | Western | 11 |
| Mbukushu | (14) | Africa | Southern Africa | SouthernAfrica_SBC | Atlantic-Congo Family | Volta-Congo | Benue-Congo | 8 |
| Kwangali | (14) | Africa | Southern Africa | SouthernAfrica_SBW | Atlantic-Congo Family | Volta-Congo | Benue-Congo | 7 |
| Mandinka | (10) | Africa | Western Africa | WesternAfrica_ME | Mande Family | Western Mande | Manding-Kpelle | 20 |
| Fula | (10) | Africa | Western Africa | WesternAfrica_NCA | Atlantic-Congo Family | North-Central Atlantic | Fula-Sereer | 20 |
| Jola | (10) | Africa | Western Africa | WesternAfrica_NCA | Atlantic-Congo Family | North-Central Atlantic | Central Atlantic | 20 |
| Wolof | (10) | Africa | Western Africa | WesternAfrica_NCA | Atlantic-Congo Family | North-Central Atlantic | Wolof-BKK | 20 |
| YRI | (13) | Africa | Western Africa | WesternAfrica_VC | Atlantic-Congo Family | Volta-Congo | Benue-Congo | 20 |
| Ga-Adangbe | (10) | Africa | Western Africa | WesternAfrica_VC | Atlantic-Congo Family | Volta-Congo | Kwa Volta-Congo | 20 |
| Igbo | (10) | Africa | Western Africa | WesternAfrica_VC | Atlantic-Congo Family | Volta-Congo | Benue-Congo | 20 |
| JPT | (13) | Asia | Eastern Asia | EasternAsia | Japonic Family | | | 20 |
| CDX | (13) | Asia | Eastern Asia | EasternAsia | Tai-Kadai Family | Kam-Tai | Be-Tai | 20 |
| CHB | (13) | Asia | Eastern Asia | EasternAsia | Sino-Tibetan Family | Sinitic | Classical-Middle-Modern Sinitic | 20 |
| KHV | (13) | Asia | South-Eastern Asia | South-EasternAsia | Austroasiatic Family | Vietic | Viet-Muong | 20 |
| GIH | (13) | Asia | Southern Asia | SouthernAsia | Indo-European Family | Indo-Iranian | Indic | 20 |
| FIN | (13) | Europe | Northern Europe | NorthernEurope | Uralic Family | Finnic | Coastal Finnic | 20 |
| GBR | (13) | Europe | Northern Europe | NorthernEurope | Indo-European Family | Germanic | Northwestern Germanic | 20 |
| IBS | (13) | Europe | Southern Europe | SouthernEurope | Indo-European Family | Italic | Romance | 20 |
| TSI | (13) | Europe | Southern Europe | SouthernEurope | Indo-European Family | Italic | Romance | 20 |

*Table_S 2: Pairwise post-hoc test results for significant differences between populations when comparing ADMIXTURE components at K=9. In the matrix of p-values, the upper right are Nemenyi post-hoc results and the lower left are Baumgartner-Weiß-Schindler results. Abbreviations; D.F. - Degrees of freedom.*

| ADMIXTURE component & proportion in modal population | Test statistic | D.F. | P-value | Population | Chrissie San | Karretjie | ‡Khomani | Nama | Zambian | Zimbabwean | BaPhuthi | AmaNdebele | AmaZulu | Duma San | BaSotho | Himba |
|---|---|---|---|---|---|---|---|---|---|---|---|---|---|---|---|---|
| BaPhuthi 0.80 | 151 | 11 | 9,39E-27 | Chrissie San | NA | 0,99 | 0,88 | 0,59 | 1,00 | 1,00 | 0,62 | 1,00 | 0,93 | 0,99 | 1,00 | 0,44 |
| | | | | Karretjie | 0,08 | NA | 1,00 | 0,87 | 1,00 | 0,91 | 0,00 | 0,17 | 0,00 | 0,09 | 0,02 | 0,74 |
| | | | | ‡Khomani | 0,03 | 0,09 | NA | 1,00 | 0,92 | 0,40 | 0,00 | 0,02 | 0,00 | 0,01 | 0,00 | 0,98 |
| | | | | Nama | 0,00 | 0,00 | 0,04 | NA | 0,50 | 0,07 | 0,00 | 0,00 | 0,00 | 0,00 | 0,00 | 1,00 |
| | | | | Zambian | 0,08 | 0,00 | 0,00 | 0,00 | NA | 1,00 | 0,00 | 0,84 | 0,03 | 0,57 | 0,69 | 0,38 |
| | | | | Zimbabwean | 0,14 | 0,00 | 0,00 | 0,00 | 0,00 | NA | 0,00 | 0,98 | 0,09 | 0,81 | 0,94 | 0,07 |
| | | | | BaPhuthi | 0,00 | 0,00 | 0,00 | 0,00 | 0,00 | 0,00 | NA | 0,65 | 1,00 | 1,00 | 0,13 | 0,00 |
| | | | | AmaNdebele | 0,04 | 0,00 | 0,00 | 0,00 | 0,00 | 0,00 | 0,00 | NA | 0,99 | 1,00 | 1,00 | 0,00 |
| | | | | AmaZulu | 0,01 | 0,00 | 0,00 | 0,00 | 0,00 | 0,00 | 0,00 | 0,00 | NA | 1,00 | 0,83 | 0,00 |
| | | | | DumaSan | 0,08 | 0,00 | 0,00 | 0,00 | 0,01 | 0,00 | 0,04 | 0,12 | 0,83 | NA | 1,00 | 0,00 |
| | | | | BaSotho | 0,02 | 0,00 | 0,00 | 0,00 | 0,00 | 0,00 | 0,00 | 0,83 | 0,00 | 0,21 | NA | 0,00 |
| | | | | Himba | 0,00 | 0,00 | 0,00 | 0,00 | 0,00 | 0,00 | 0,00 | 0,00 | 0,00 | 0,00 | 0,00 | NA |
| GBR 0.99 | 95 | 11 | 2,07E-15 | Chrissie San | NA | 0,60 | 0,46 | 0,35 | 1,00 | 1,00 | 1,00 | 1,00 | 1,00 | 1,00 | 1,00 | 1,00 |
| | | | | Karretjie | 0,00 | NA | 1,00 | 1,00 | 0,06 | 0,05 | 0,00 | 0,08 | 0,05 | 0,61 | 0,02 | 0,08 |
| | | | | ‡Khomani | 0,00 | 0,05 | NA | 1,00 | 0,02 | 0,01 | 0,00 | 0,03 | 0,01 | 0,42 | 0,00 | 0,03 |
| | | | | Nama | 0,00 | 0,35 | 0,35 | NA | 0,01 | 0,01 | 0,00 | 0,01 | 0,00 | 0,30 | 0,00 | 0,01 |
| | | | | Zambian | 0,00 | 0,00 | 0,00 | 0,00 | NA | 1,00 | 1,00 | 1,00 | 1,00 | 1,00 | 1,00 | 1,00 |
| | | | | Zimbabwean | 0,00 | 0,00 | 0,00 | 0,00 | 0,00 | NA | 1,00 | 1,00 | 1,00 | 1,00 | 1,00 | 1,00 |
| | | | | BaPhuthi | 0,00 | 0,00 | 0,00 | 0,00 | 0,00 | 0,00 | NA | 1,00 | 1,00 | 1,00 | 1,00 | 1,00 |
| | | | | AmaNdebele | 0,00 | 0,00 | 0,00 | 0,00 | 0,00 | 0,00 | 0,00 | NA | 1,00 | 1,00 | 1,00 | 1,00 |
| | | | | AmaZulu | 0,00 | 0,00 | 0,00 | 0,00 | 0,00 | 0,00 | 0,00 | 0,00 | NA | 1,00 | 1,00 | 1,00 |
| | | | | DumaSan | 0,00 | 0,00 | 0,00 | 0,00 | 0,00 | 0,00 | 0,00 | 0,00 | 0,00 | NA | 1,00 | 1,00 |

| | | | | | | | | | | | | | | | | |
|---|---|---|---|---|---|---|---|---|---|---|---|---|---|---|---|---|
| | | | | BaSotho | 0,00 | 0,00 | 0,00 | 0,00 | 0,00 | 0,00 | 0,00 | 0,00 | 0,00 | 0,00 | NA | 1,00 |
| | | | | Himba | 0,00 | 0,00 | 0,00 | 0,00 | 0,00 | 0,00 | 0,00 | 0,00 | 0,00 | 0,00 | 0,00 | NA |
| Himba 0.92 | 119 | 11 | 2,36E-20 | Chrissie San | NA | 1,00 | 1,00 | 0,53 | 0,00 | 0,01 | 1,00 | 0,37 | 0,64 | 0,94 | 0,14 | 0,00 |
| | | | | Karretjie | 0,00 | NA | 1,00 | 0,00 | 0,00 | 0,00 | 0,73 | 0,01 | 0,01 | 0,64 | 0,00 | 0,00 |
| | | | | ‡Khomani | 0,00 | 0,00 | NA | 0,06 | 0,00 | 0,00 | 1,00 | 0,08 | 0,14 | 0,95 | 0,00 | 0,00 |
| | | | | Nama | 0,00 | 0,00 | 0,00 | NA | 0,07 | 0,19 | 0,43 | 1,00 | 1,00 | 1,00 | 0,98 | 0,02 |
| | | | | Zambian | 0,06 | 0,00 | 0,00 | 0,01 | NA | 1,00 | 0,00 | 0,67 | 0,03 | 0,18 | 0,59 | 1,00 |
| | | | | Zimbabwean | 0,02 | 0,00 | 0,00 | 0,01 | 0,00 | NA | 0,00 | 0,90 | 0,09 | 0,36 | 0,88 | 1,00 |
| | | | | BaPhuthi | 0,00 | 0,00 | 0,00 | 0,00 | 0,00 | 0,00 | NA | 0,37 | 0,64 | 1,00 | 0,01 | 0,00 |
| | | | | AmaNdebele | 0,06 | 0,00 | 0,00 | 0,09 | 0,00 | 0,00 | 0,00 | NA | 1,00 | 1,00 | 1,00 | 0,36 |
| | | | | AmaZulu | 0,01 | 0,00 | 0,00 | 0,09 | 0,00 | 0,00 | 0,00 | 0,07 | NA | 1,00 | 0,91 | 0,01 |
| | | | | DumaSan | 0,09 | 0,00 | 0,00 | 0,09 | 0,01 | 0,00 | 0,09 | 0,14 | 0,20 | NA | 0,96 | 0,07 |
| | | | | BaSotho | 0,01 | 0,00 | 0,00 | 0,06 | 0,00 | 0,00 | 0,00 | 0,14 | 0,00 | 0,06 | NA | 0,26 |
| | | | | Himba | 0,06 | 0,00 | 0,00 | 0,00 | 0,00 | 0,00 | 0,00 | 0,00 | 0,00 | 0,01 | 0,00 | NA |
| CDX 1.00 | 76 | 11 | 7,20E-12 | Chrissie San | NA | 0,34 | 0,82 | 1,00 | 1,00 | 1,00 | 1,00 | 1,00 | 1,00 | 1,00 | 1,00 | 1,00 |
| | | | | Karretjie | 0,00 | NA | 0,97 | 0,12 | 0,01 | 0,00 | 0,00 | 0,01 | 0,00 | 0,82 | 0,00 | 0,01 |
| | | | | ‡Khomani | 0,00 | 0,02 | NA | 0,89 | 0,19 | 0,09 | 0,03 | 0,24 | 0,18 | 1,00 | 0,10 | 0,24 |
| | | | | Nama | 0,00 | 0,00 | 0,00 | NA | 0,95 | 0,92 | 0,87 | 0,97 | 0,99 | 1,00 | 0,98 | 0,97 |
| | | | | Zambian | 0,00 | 0,00 | 0,00 | 0,00 | NA | 1,00 | 1,00 | 1,00 | 1,00 | 0,99 | 1,00 | 1,00 |
| | | | | Zimbabwean | 0,00 | 0,00 | 0,00 | 0,00 | 0,00 | NA | 1,00 | 1,00 | 1,00 | 0,99 | 1,00 | 1,00 |
| | | | | BaPhuthi | 0,00 | 0,00 | 0,00 | 0,00 | 0,00 | 0,00 | NA | 1,00 | 1,00 | 0,99 | 1,00 | 1,00 |
| | | | | AmaNdebele | 0,00 | 0,00 | 0,00 | 0,00 | 0,00 | 0,00 | 0,00 | NA | 1,00 | 0,99 | 1,00 | 1,00 |
| | | | | AmaZulu | 0,00 | 0,00 | 0,00 | 0,00 | 0,00 | 0,00 | 0,00 | 0,00 | NA | 1,00 | 1,00 | 1,00 |
| | | | | DumaSan | 0,01 | 0,02 | 0,00 | 0,00 | 0,00 | 0,00 | 0,00 | 0,00 | 0,00 | NA | 1,00 | 0,99 |
| | | | | BaSotho | 0,00 | 0,00 | 0,00 | 0,00 | 0,00 | 0,00 | 0,00 | 0,00 | 0,00 | 0,00 | NA | 1,00 |
| | | | | Himba | 0,00 | 0,00 | 0,00 | 0,00 | 0,00 | 0,00 | 0,00 | 0,00 | 0,00 | 0,00 | 0,00 | NA |
| Somali 0.92 | 105 | 11 | 1,52E-17 | Chrissie San | NA | 1,00 | 0,16 | 0,01 | 1,00 | 1,00 | 0,99 | 1,00 | 0,49 | 0,59 | 1,00 | 1,00 |
| | | | | Karretjie | 0,00 | NA | 0,01 | 0,00 | 0,99 | 0,98 | 1,00 | 1,00 | 0,17 | 0,59 | 1,00 | 1,00 |
| | | | | ‡Khomani | 0,00 | 0,00 | NA | 0,75 | 0,00 | 0,00 | 0,04 | 0,04 | 0,99 | 1,00 | 0,00 | 0,07 |
| | | | | Nama | 0,00 | 0,00 | 0,00 | NA | 0,00 | 0,00 | 0,00 | 0,00 | 0,10 | 0,90 | 0,00 | 0,00 |
| | | | | Zambian | 0,00 | 0,00 | 0,00 | 0,00 | NA | 1,00 | 0,77 | 1,00 | 0,02 | 0,18 | 1,00 | 1,00 |

| | | | | | | | | | | | | | | | | | |
|---|---|---|---|---|---|---|---|---|---|---|---|---|---|---|---|---|---|
| | | | | | Zimbabwean | 0,00 | 0,00 | 0,00 | 0,00 | 0,00 | NA | 0,64 | 1,00 | 0,01 | 0,13 | 1,00 | 1,00 |
| | | | | | BaPhuthi | 0,00 | 0,00 | 0,00 | 0,00 | 0,00 | 0,00 | NA | 1,00 | 0,55 | 0,88 | 0,99 | 1,00 |
| | | | | | AmaNdebele | 0,00 | 0,00 | 0,00 | 0,00 | 0,00 | 0,00 | 0,00 | NA | 0,36 | 0,63 | 1,00 | 1,00 |
| | | | | | AmaZulu | 0,00 | 0,00 | 0,00 | 0,00 | 0,00 | 0,00 | 0,00 | 0,00 | NA | 1,00 | 0,05 | 0,46 |
| | | | | | DumaSan | 0,06 | 0,00 | 0,06 | 0,00 | 0,00 | 0,00 | 0,00 | 0,02 | 0,87 | NA | 0,42 | 0,71 |
| | | | | | BaSotho | 0,00 | 0,00 | 0,00 | 0,00 | 0,00 | 0,00 | 0,00 | 0,00 | 0,00 | 0,00 | NA | 1,00 |
| | | | | | Himba | 0,00 | 0,00 | 0,00 | 0,00 | 0,00 | 0,00 | 0,00 | 0,00 | 0,00 | 0,02 | 0,00 | NA |
| Jola | 0.98 | 106 | 11 | 1,01E-17 | Chrissie San | NA | 1,00 | 1,00 | 1,00 | 0,32 | 0,45 | 1,00 | 1,00 | 1,00 | 1,00 | 0,98 | 1,00 |
| | | | | | Karretjie | 0,00 | NA | 1,00 | 1,00 | 0,00 | 0,00 | 0,98 | 0,68 | 0,51 | 0,22 | 0,00 | 1,00 |
| | | | | | ‡Khomani | 0,00 | 0,00 | NA | 1,00 | 0,00 | 0,00 | 0,97 | 0,65 | 0,44 | 0,20 | 0,00 | 1,00 |
| | | | | | Nama | 0,00 | 0,00 | 0,00 | NA | 0,00 | 0,00 | 1,00 | 0,98 | 0,98 | 0,63 | 0,03 | 1,00 |
| | | | | | Zambian | 0,02 | 0,00 | 0,00 | 0,00 | NA | 1,00 | 0,00 | 0,11 | 0,00 | 0,87 | 0,53 | 0,00 |
| | | | | | Zimbabwean | 0,03 | 0,00 | 0,00 | 0,00 | 0,18 | NA | 0,00 | 0,18 | 0,01 | 0,96 | 0,73 | 0,00 |
| | | | | | BaPhuthi | 0,00 | 0,00 | 0,00 | 0,00 | 0,00 | 0,00 | NA | 0,99 | 0,99 | 0,71 | 0,04 | 1,00 |
| | | | | | AmaNdebele | 0,15 | 0,00 | 0,00 | 0,00 | 0,00 | 0,00 | 0,00 | NA | 1,00 | 1,00 | 0,97 | 0,95 |
| | | | | | AmaZulu | 0,00 | 0,00 | 0,00 | 0,00 | 0,00 | 0,00 | 0,00 | 0,00 | NA | 0,98 | 0,57 | 0,95 |
| | | | | | DumaSan | 0,61 | 0,00 | 0,00 | 0,00 | 0,00 | 0,01 | 0,00 | 0,61 | 0,00 | NA | 1,00 | 0,55 |
| | | | | | BaSotho | 0,01 | 0,00 | 0,00 | 0,00 | 0,00 | 0,00 | 0,00 | 0,01 | 0,00 | 0,18 | NA | 0,08 |
| | | | | | Himba | 0,00 | 0,00 | 0,00 | 0,00 | 0,00 | 0,00 | 0,00 | 0,00 | 0,00 | 0,00 | 0,00 | NA |
| GIH | 0.93 | 88 | 11 | 3,46E-14 | Chrissie San | NA | 0,44 | 0,54 | 0,92 | 1,00 | 1,00 | 1,00 | 1,00 | 1,00 | 1,00 | 1,00 | 1,00 |
| | | | | | Karretjie | 0,00 | NA | 1,00 | 0,94 | 0,02 | 0,01 | 0,00 | 0,09 | 0,00 | 0,54 | 0,00 | 0,03 |
| | | | | | ‡Khomani | 0,00 | 0,01 | NA | 0,98 | 0,03 | 0,01 | 0,01 | 0,15 | 0,00 | 0,66 | 0,00 | 0,05 |
| | | | | | Nama | 0,00 | 0,00 | 0,00 | NA | 0,41 | 0,26 | 0,32 | 0,75 | 0,17 | 0,99 | 0,10 | 0,48 |
| | | | | | Zambian | 0,00 | 0,00 | 0,00 | 0,00 | NA | 1,00 | 1,00 | 1,00 | 1,00 | 1,00 | 1,00 | 1,00 |
| | | | | | Zimbabwean | 0,00 | 0,00 | 0,00 | 0,00 | 0,00 | NA | 1,00 | 1,00 | 1,00 | 1,00 | 1,00 | 1,00 |
| | | | | | BaPhuthi | 0,00 | 0,00 | 0,00 | 0,00 | 0,00 | 0,00 | NA | 1,00 | 1,00 | 1,00 | 1,00 | 1,00 |
| | | | | | AmaNdebele | 0,00 | 0,00 | 0,00 | 0,00 | 0,00 | 0,00 | 0,00 | NA | 1,00 | 1,00 | 1,00 | 1,00 |
| | | | | | AmaZulu | 0,00 | 0,00 | 0,00 | 0,00 | 0,00 | 0,00 | 0,00 | 0,00 | NA | 1,00 | 1,00 | 1,00 |
| | | | | | DumaSan | 0,00 | 0,00 | 0,00 | 0,00 | 0,00 | 0,00 | 0,00 | 0,00 | 0,00 | NA | 1,00 | 1,00 |
| | | | | | BaSotho | 0,00 | 0,00 | 0,00 | 0,00 | 0,00 | 0,00 | 0,00 | 0,00 | 0,00 | 0,00 | NA | 1,00 |
| | | | | | Himba | 0,00 | 0,00 | 0,00 | 0,00 | 0,00 | 0,00 | 0,00 | 0,00 | 0,00 | 0,00 | 0,00 | NA |
| Naro | 0.98 | 142 | 11 | 7,16E- | Chrissie San | NA | 1,00 | 1,00 | 1,00 | 0,06 | 0,02 | 0,93 | 1,00 | 0,57 | 0,99 | 0,90 | 0,18 |

| | | | | | | | | | | | | | | | | | |
|---|---|---|---|---|---|---|---|---|---|---|---|---|---|---|---|---|---|
| | | | | 25 | | | | | | | | | | | | | |
| | | | | | Karretjie | 0,11 | NA | 1,00 | 1,00 | 0,00 | 0,00 | 0,00 | 0,25 | 0,00 | 0,23 | 0,00 | 0,00 |
| | | | | | ‡Khomani | 0,32 | 1,00 | NA | 1,00 | 0,00 | 0,00 | 0,00 | 0,41 | 0,00 | 0,36 | 0,00 | 0,00 |
| | | | | | Nama | 0,91 | 0,91 | 1,00 | NA | 0,00 | 0,00 | 0,00 | 0,58 | 0,00 | 0,50 | 0,00 | 0,00 |
| | | | | | Zambian | 0,13 | 0,00 | 0,00 | 0,00 | NA | 1,00 | 0,09 | 0,06 | 0,68 | 0,44 | 0,15 | 1,00 |
| | | | | | Zimbabwean | 0,00 | 0,00 | 0,00 | 0,00 | 0,04 | NA | 0,01 | 0,01 | 0,32 | 0,24 | 0,03 | 1,00 |
| | | | | | BaPhuthi | 0,04 | 0,00 | 0,00 | 0,00 | 0,00 | 0,00 | NA | 1,00 | 0,98 | 1,00 | 1,00 | 0,48 |
| | | | | | AmaNdebele | 0,15 | 0,00 | 0,00 | 0,00 | 0,00 | 0,00 | 0,42 | NA | 0,84 | 1,00 | 1,00 | 0,30 |
| | | | | | AmaZulu | 0,05 | 0,00 | 0,00 | 0,00 | 0,00 | 0,00 | 0,02 | 0,00 | NA | 1,00 | 1,00 | 0,98 |
| | | | | | DumaSan | 0,25 | 0,00 | 0,00 | 0,00 | 0,01 | 0,00 | 1,00 | 1,00 | 0,13 | NA | 1,00 | 0,79 |
| | | | | | BaSotho | 0,05 | 0,00 | 0,00 | 0,00 | 0,00 | 0,00 | 1,00 | 0,91 | 0,20 | 1,00 | NA | 0,61 |
| | | | | | Himba | 0,15 | 0,00 | 0,00 | 0,00 | 0,00 | 0,00 | 0,00 | 0,00 | 0,00 | 0,02 | 0,00 | NA |
| Kalenjin | 0.68 | 102 | 11 | 7,00E-17 | Chrissie San | NA | 1,00 | 1,00 | 1,00 | 0,02 | 0,02 | 1,00 | 0,87 | 0,93 | 0,83 | 0,62 | 1,00 |
| | | | | | Karretjie | 0,00 | NA | 1,00 | 1,00 | 0,00 | 0,00 | 1,00 | 0,34 | 0,25 | 0,40 | 0,01 | 1,00 |
| | | | | | ‡Khomani | 0,00 | 0,00 | NA | 1,00 | 0,00 | 0,00 | 1,00 | 0,37 | 0,26 | 0,43 | 0,01 | 1,00 |
| | | | | | Nama | 0,00 | 0,00 | 0,00 | NA | 0,00 | 0,00 | 1,00 | 0,88 | 0,91 | 0,86 | 0,22 | 1,00 |
| | | | | | Zambian | 0,03 | 0,00 | 0,00 | 0,00 | NA | 1,00 | 0,00 | 0,39 | 0,03 | 0,80 | 0,32 | 0,00 |
| | | | | | Zimbabwean | 0,02 | 0,00 | 0,00 | 0,00 | 0,92 | NA | 0,00 | 0,34 | 0,01 | 0,79 | 0,23 | 0,00 |
| | | | | | BaPhuthi | 0,00 | 0,00 | 0,00 | 0,00 | 0,00 | 0,00 | NA | 0,84 | 0,86 | 0,83 | 0,15 | 1,00 |
| | | | | | AmaNdebele | 0,04 | 0,00 | 0,00 | 0,00 | 0,00 | 0,00 | 0,00 | NA | 1,00 | 1,00 | 1,00 | 0,48 |
| | | | | | AmaZulu | 0,00 | 0,00 | 0,00 | 0,00 | 0,00 | 0,00 | 0,00 | 0,02 | NA | 1,00 | 0,99 | 0,49 |
| | | | | | DumaSan | 0,22 | 0,00 | 0,00 | 0,00 | 0,00 | 0,00 | 0,00 | 0,92 | 0,00 | NA | 1,00 | 0,48 |
| | | | | | BaSotho | 0,00 | 0,00 | 0,00 | 0,00 | 0,00 | 0,00 | 0,00 | 0,42 | 0,02 | 0,42 | NA | 0,08 |
| | | | | | Himba | 0,00 | 0,00 | 0,00 | 0,00 | 0,00 | 0,00 | 0,00 | 0,00 | 0,00 | 0,00 | 0,00 | NA |

*Table_S 3: Estimates of the ADMIXTURE components in each population at K = 9. Indicated are the mean ($\mu$), minimum (Min), maximim (Max) and standard deviation ($\sigma$) within each population at K = 9 for each component and for selected combinations of components.*

| ADMIXTURE components | 'BaPhuthi' | | | | 'GBR' | | | | 'Himba' | | | | 'CDX' | | | | 'Somali' | | | | 'Jola' | | | |
|---|---|---|---|---|---|---|---|---|---|---|---|---|---|---|---|---|---|---|---|---|---|---|---|---|
| Population | μ | Max | Min | σ | μ | Max | Min | σ | μ | Max | Min | σ | μ | Max | Min | σ | μ | Max | Min | σ | μ | Max | Min | σ |
| !Xun | 0.000 | 0.000 | 0.000 | 0.000 | 0.000 | 0.000 | 0.000 | 0.000 | 0.199 | 0.273 | 0.114 | 0.047 | 0.000 | 0.003 | 0.000 | 0.001 | 0.000 | 0.009 | 0.000 | 0.002 | 0.008 | 0.053 | 0.000 | 0.016 |
| CDX | 0.000 | 0.000 | 0.000 | 0.000 | 0.000 | 0.000 | 0.000 | 0.000 | 0.000 | 0.000 | 0.000 | 0.000 | 1.000 | 1.000 | 1.000 | 0.000 | 0.000 | 0.000 | 0.000 | 0.000 | 0.000 | 0.000 | 0.000 | 0.000 |
| CHB | 0.000 | 0.000 | 0.000 | 0.000 | 0.003 | 0.022 | 0.000 | 0.007 | 0.000 | 0.000 | 0.000 | 0.000 | 0.995 | 1.000 | 0.967 | 0.010 | 0.000 | 0.000 | 0.000 | 0.000 | 0.000 | 0.000 | 0.000 | 0.000 |
| FIN | 0.000 | 0.000 | 0.000 | 0.000 | 0.933 | 0.957 | 0.910 | 0.011 | 0.000 | 0.000 | 0.000 | 0.000 | 0.064 | 0.090 | 0.034 | 0.013 | 0.000 | 0.000 | 0.000 | 0.000 | 0.000 | 0.000 | 0.000 | 0.000 |
| IBS | 0.000 | 0.000 | 0.000 | 0.000 | 0.923 | 0.972 | 0.863 | 0.029 | 0.000 | 0.000 | 0.000 | 0.000 | 0.001 | 0.009 | 0.000 | 0.002 | 0.074 | 0.137 | 0.028 | 0.027 | 0.001 | 0.011 | 0.000 | 0.003 |
| KHV | 0.000 | 0.000 | 0.000 | 0.000 | 0.000 | 0.007 | 0.000 | 0.002 | 0.000 | 0.000 | 0.000 | 0.000 | 0.990 | 1.000 | 0.970 | 0.011 | 0.000 | 0.000 | 0.000 | 0.000 | 0.000 | 0.000 | 0.000 | 0.000 |
| Naro | 0.000 | 0.000 | 0.000 | 0.000 | 0.000 | 0.000 | 0.000 | 0.000 | 0.018 | 0.025 | 0.011 | 0.010 | 0.000 | 0.000 | 0.000 | 0.000 | 0.000 | 0.000 | 0.000 | 0.000 | 0.000 | 0.000 | 0.000 | 0.000 |
| TSI | 0.000 | 0.000 | 0.000 | 0.000 | 0.912 | 0.929 | 0.890 | 0.011 | 0.000 | 0.000 | 0.000 | 0.000 | 0.000 | 0.000 | 0.000 | 0.000 | 0.069 | 0.087 | 0.056 | 0.007 | 0.000 | 0.000 | 0.000 | 0.000 |
| Ju\|'hoansi | 0.005 | 0.073 | 0.000 | 0.020 | 0.000 | 0.000 | 0.000 | 0.000 | 0.020 | 0.284 | 0.000 | 0.076 | 0.000 | 0.000 | 0.000 | 0.000 | 0.000 | 0.000 | 0.000 | 0.000 | 0.002 | 0.031 | 0.000 | 0.008 |
| Hai\|\|om | 0.019 | 0.058 | 0.000 | 0.024 | 0.004 | 0.017 | 0.000 | 0.007 | 0.296 | 0.346 | 0.233 | 0.042 | 0.001 | 0.003 | 0.000 | 0.001 | 0.119 | 0.147 | 0.096 | 0.016 | 0.009 | 0.025 | 0.000 | 0.010 |
| Nama | 0.031 | 0.178 | 0.000 | 0.042 | 0.092 | 0.300 | 0.000 | 0.111 | 0.167 | 0.455 | 0.000 | 0.174 | 0.003 | 0.015 | 0.000 | 0.005 | 0.101 | 0.182 | 0.018 | 0.046 | 0.003 | 0.028 | 0.000 | 0.007 |
| Jola | 0.000 | 0.005 | 0.000 | 0.001 | 0.002 | 0.035 | 0.000 | 0.008 | 0.006 | 0.113 | 0.000 | 0.025 | 0.000 | 0.002 | 0.000 | 0.000 | 0.002 | 0.034 | 0.000 | 0.008 | 0.981 | 1.000 | 0.784 | 0.048 |
| Khwe | 0.042 | 0.111 | 0.000 | 0.038 | 0.001 | 0.005 | 0.000 | 0.001 | 0.507 | 0.609 | 0.427 | 0.062 | 0.001 | 0.007 | 0.000 | 0.002 | 0.052 | 0.082 | 0.023 | 0.017 | 0.015 | 0.049 | 0.000 | 0.019 |
| ‡Khomani | 0.101 | 0.408 | 0.000 | 0.110 | 0.120 | 0.408 | 0.000 | 0.134 | 0.023 | 0.212 | 0.000 | 0.052 | 0.017 | 0.077 | 0.000 | 0.023 | 0.048 | 0.111 | 0.000 | 0.035 | 0.000 | 0.000 | 0.000 | 0.000 |
| Oromo | 0.002 | 0.019 | 0.000 | 0.004 | 0.150 | 0.194 | 0.108 | 0.021 | 0.001 | 0.014 | 0.000 | 0.004 | 0.003 | 0.015 | 0.000 | 0.005 | 0.706 | 0.743 | 0.667 | 0.022 | 0.003 | 0.015 | 0.000 | 0.005 |
| Chrissie San | 0.538 | 0.567 | 0.514 | 0.027 | 0.000 | 0.000 | 0.000 | 0.000 | 0.000 | 0.000 | 0.000 | 0.000 | 0.000 | 0.000 | 0.000 | 0.000 | 0.000 | 0.000 | 0.000 | 0.000 | 0.013 | 0.039 | 0.000 | 0.023 |
| Himba | 0.006 | 0.044 | 0.000 | 0.016 | 0.000 | 0.000 | 0.000 | 0.000 | 0.922 | 0.953 | 0.864 | 0.027 | 0.000 | 0.000 | 0.000 | 0.000 | 0.004 | 0.018 | 0.000 | 0.006 | 0.000 | 0.001 | 0.000 | 0.000 |
| Karretjie | 0.177 | 0.531 | 0.014 | 0.119 | 0.058 | 0.213 | 0.000 | 0.063 | 0.001 | 0.012 | 0.000 | 0.003 | 0.016 | 0.063 | 0.000 | 0.016 | 0.006 | 0.040 | 0.000 | 0.012 | 0.000 | 0.000 | 0.000 | 0.000 |
| G\|ui & G\|\|ana | 0.290 | 0.502 | 0.106 | 0.128 | 0.000 | 0.000 | 0.000 | 0.000 | 0.012 | 0.029 | 0.000 | 0.013 | 0.000 | 0.000 | 0.000 | 0.000 | 0.004 | 0.026 | 0.000 | 0.008 | 0.004 | 0.020 | 0.000 | 0.007 |
| BaPhuthi_SA | 0.768 | 0.784 | 0.752 | 0.023 | 0.000 | 0.000 | 0.000 | 0.000 | 0.043 | 0.063 | 0.024 | 0.027 | 0.000 | 0.000 | 0.000 | 0.000 | 0.013 | 0.026 | 0.000 | 0.018 | 0.001 | 0.002 | 0.000 | 0.002 |
| Mbukushu | 0.247 | 0.274 | 0.218 | 0.021 | 0.000 | 0.001 | 0.000 | 0.000 | 0.537 | 0.616 | 0.481 | 0.040 | 0.001 | 0.004 | 0.000 | 0.001 | 0.001 | 0.006 | 0.000 | 0.002 | 0.087 | 0.104 | 0.041 | 0.021 |
| AmaNdebele | 0.699 | 0.735 | 0.663 | 0.024 | 0.000 | 0.000 | 0.000 | 0.000 | 0.089 | 0.103 | 0.076 | 0.010 | 0.000 | 0.000 | 0.000 | 0.000 | 0.005 | 0.029 | 0.000 | 0.010 | 0.010 | 0.029 | 0.000 | 0.011 |
| BaPhuthi_LE | 0.806 | 0.872 | 0.691 | 0.049 | 0.001 | 0.012 | 0.000 | 0.003 | 0.028 | 0.095 | 0.000 | 0.030 | 0.000 | 0.001 | 0.000 | 0.000 | 0.008 | 0.046 | 0.000 | 0.012 | 0.004 | 0.031 | 0.000 | 0.009 |
| AmaZulu | 0.757 | 0.796 | 0.710 | 0.022 | 0.002 | 0.026 | 0.000 | 0.006 | 0.070 | 0.118 | 0.034 | 0.021 | 0.001 | 0.026 | 0.000 | 0.006 | 0.019 | 0.040 | 0.000 | 0.011 | 0.010 | 0.034 | 0.000 | 0.013 |
| DumaSan | 0.723 | 0.781 | 0.577 | 0.084 | 0.000 | 0.002 | 0.000 | 0.001 | 0.053 | 0.093 | 0.004 | 0.035 | 0.004 | 0.012 | 0.000 | 0.005 | 0.021 | 0.032 | 0.003 | 0.012 | 0.015 | 0.033 | 0.003 | 0.014 |
| Damara | 0.024 | 0.113 | 0.000 | 0.040 | 0.000 | 0.000 | 0.000 | 0.000 | 0.803 | 0.932 | 0.689 | 0.092 | 0.000 | 0.001 | 0.000 | 0.000 | 0.023 | 0.062 | 0.000 | 0.026 | 0.006 | 0.034 | 0.000 | 0.012 |
| BaSotho | 0.681 | 0.755 | 0.564 | 0.050 | 0.005 | 0.075 | 0.000 | 0.018 | 0.115 | 0.200 | 0.032 | 0.043 | 0.001 | 0.006 | 0.000 | 0.002 | 0.003 | 0.014 | 0.000 | 0.004 | 0.029 | 0.056 | 0.000 | 0.020 |
| Amhara | 0.001 | 0.013 | 0.000 | 0.003 | 0.189 | 0.212 | 0.155 | 0.017 | 0.001 | 0.009 | 0.000 | 0.002 | 0.001 | 0.006 | 0.000 | 0.002 | 0.716 | 0.746 | 0.696 | 0.014 | 0.004 | 0.026 | 0.000 | 0.008 |
| Kwangali | 0.255 | 0.293 | 0.231 | 0.024 | 0.000 | 0.000 | 0.000 | 0.000 | 0.538 | 0.589 | 0.518 | 0.027 | 0.000 | 0.000 | 0.000 | 0.000 | 0.001 | 0.004 | 0.000 | 0.001 | 0.101 | 0.156 | 0.078 | 0.025 |
| Kongo | 0.269 | 0.280 | 0.253 | 0.012 | 0.000 | 0.000 | 0.000 | 0.000 | 0.468 | 0.497 | 0.437 | 0.025 | 0.000 | 0.000 | 0.000 | 0.000 | 0.000 | 0.000 | 0.000 | 0.000 | 0.149 | 0.176 | 0.115 | 0.023 |
| JPT | 0.000 | 0.000 | 0.000 | 0.000 | 0.000 | 0.003 | 0.000 | 0.001 | 0.000 | 0.000 | 0.000 | 0.000 | 0.997 | 1.000 | 0.986 | 0.005 | 0.000 | 0.000 | 0.000 | 0.000 | 0.000 | 0.000 | 0.000 | 0.000 |

| | | | | | | | | | | | | | | | | | | | | | | | | |
|---|---|---|---|---|---|---|---|---|---|---|---|---|---|---|---|---|---|---|---|---|---|---|---|---|
| GBR | 0.000 | 0.001 | 0.000 | 0.000 | 0.989 | 1.000 | 0.969 | 0.008 | 0.001 | 0.008 | 0.000 | 0.002 | 0.001 | 0.005 | 0.000 | 0.002 | 0.002 | 0.018 | 0.000 | 0.005 | 0.000 | 0.003 | 0.000 | 0.001 |
| GIH | 0.000 | 0.007 | 0.000 | 0.001 | 0.061 | 0.328 | 0.000 | 0.112 | 0.000 | 0.005 | 0.000 | 0.001 | 0.011 | 0.043 | 0.000 | 0.015 | 0.000 | 0.006 | 0.000 | 0.001 | 0.000 | 0.000 | 0.000 | 0.000 |
| MKK | 0.012 | 0.060 | 0.000 | 0.019 | 0.008 | 0.032 | 0.000 | 0.010 | 0.006 | 0.045 | 0.000 | 0.012 | 0.001 | 0.006 | 0.000 | 0.001 | 0.420 | 0.452 | 0.365 | 0.026 | 0.001 | 0.015 | 0.000 | 0.004 |
| Mandinka | 0.019 | 0.069 | 0.000 | 0.025 | 0.003 | 0.025 | 0.000 | 0.006 | 0.055 | 0.177 | 0.000 | 0.049 | 0.000 | 0.004 | 0.000 | 0.001 | 0.006 | 0.021 | 0.000 | 0.007 | 0.892 | 0.996 | 0.738 | 0.071 |
| Kalenjin | 0.011 | 0.063 | 0.000 | 0.017 | 0.000 | 0.000 | 0.000 | 0.000 | 0.008 | 0.058 | 0.000 | 0.015 | 0.000 | 0.001 | 0.000 | 0.000 | 0.283 | 0.322 | 0.197 | 0.033 | 0.013 | 0.075 | 0.000 | 0.020 |
| Fula | 0.016 | 0.081 | 0.000 | 0.026 | 0.075 | 0.183 | 0.000 | 0.066 | 0.051 | 0.189 | 0.000 | 0.066 | 0.000 | 0.004 | 0.000 | 0.001 | 0.072 | 0.154 | 0.000 | 0.061 | 0.765 | 0.943 | 0.660 | 0.074 |
| Somali | 0.018 | 0.070 | 0.000 | 0.022 | 0.011 | 0.154 | 0.000 | 0.035 | 0.020 | 0.056 | 0.000 | 0.015 | 0.000 | 0.002 | 0.000 | 0.000 | 0.924 | 0.976 | 0.691 | 0.072 | 0.017 | 0.052 | 0.000 | 0.015 |
| YRI | 0.155 | 0.196 | 0.129 | 0.019 | 0.000 | 0.000 | 0.000 | 0.000 | 0.397 | 0.450 | 0.346 | 0.026 | 0.000 | 0.002 | 0.000 | 0.000 | 0.001 | 0.022 | 0.000 | 0.005 | 0.425 | 0.478 | 0.368 | 0.027 |
| Wolof | 0.019 | 0.089 | 0.000 | 0.024 | 0.005 | 0.015 | 0.000 | 0.005 | 0.033 | 0.149 | 0.000 | 0.040 | 0.001 | 0.005 | 0.000 | 0.002 | 0.028 | 0.050 | 0.001 | 0.015 | 0.893 | 0.977 | 0.775 | 0.047 |
| Ga-Adangbe | 0.122 | 0.157 | 0.081 | 0.021 | 0.001 | 0.021 | 0.000 | 0.005 | 0.368 | 0.422 | 0.314 | 0.032 | 0.001 | 0.005 | 0.000 | 0.001 | 0.001 | 0.015 | 0.000 | 0.004 | 0.491 | 0.557 | 0.452 | 0.026 |
| Kikuyu | 0.185 | 0.233 | 0.145 | 0.027 | 0.021 | 0.034 | 0.000 | 0.009 | 0.122 | 0.192 | 0.079 | 0.028 | 0.001 | 0.007 | 0.000 | 0.002 | 0.244 | 0.287 | 0.167 | 0.029 | 0.025 | 0.069 | 0.000 | 0.019 |
| Igbo | 0.173 | 0.205 | 0.128 | 0.023 | 0.000 | 0.000 | 0.000 | 0.000 | 0.419 | 0.477 | 0.381 | 0.026 | 0.000 | 0.007 | 0.000 | 0.001 | 0.000 | 0.006 | 0.000 | 0.001 | 0.372 | 0.407 | 0.311 | 0.023 |
| LWK | 0.237 | 0.293 | 0.187 | 0.026 | 0.001 | 0.008 | 0.000 | 0.002 | 0.243 | 0.300 | 0.177 | 0.026 | 0.000 | 0.002 | 0.000 | 0.000 | 0.006 | 0.047 | 0.000 | 0.012 | 0.038 | 0.062 | 0.007 | 0.017 |
| Barundi | 0.224 | 0.277 | 0.160 | 0.032 | 0.014 | 0.028 | 0.000 | 0.009 | 0.285 | 0.351 | 0.202 | 0.033 | 0.001 | 0.008 | 0.000 | 0.002 | 0.056 | 0.151 | 0.000 | 0.036 | 0.060 | 0.103 | 0.013 | 0.022 |
| Banyarwanda | 0.192 | 0.258 | 0.095 | 0.042 | 0.024 | 0.042 | 0.000 | 0.010 | 0.230 | 0.306 | 0.147 | 0.044 | 0.001 | 0.007 | 0.000 | 0.002 | 0.131 | 0.283 | 0.052 | 0.066 | 0.048 | 0.091 | 0.000 | 0.026 |
| Angolan | 0.259 | 0.306 | 0.232 | 0.027 | 0.000 | 0.000 | 0.000 | 0.000 | 0.510 | 0.549 | 0.457 | 0.034 | 0.000 | 0.000 | 0.000 | 0.000 | 0.000 | 0.000 | 0.000 | 0.000 | 0.132 | 0.148 | 0.111 | 0.012 |
| Yansi | 0.270 | 0.293 | 0.241 | 0.022 | 0.000 | 0.000 | 0.000 | 0.000 | 0.460 | 0.481 | 0.433 | 0.020 | 0.000 | 0.000 | 0.000 | 0.000 | 0.000 | 0.000 | 0.000 | 0.000 | 0.162 | 0.167 | 0.149 | 0.008 |
| Zambian | 0.370 | 0.425 | 0.324 | 0.039 | 0.000 | 0.000 | 0.000 | 0.000 | 0.407 | 0.456 | 0.349 | 0.035 | 0.000 | 0.000 | 0.000 | 0.000 | 0.000 | 0.000 | 0.000 | 0.000 | 0.086 | 0.121 | 0.062 | 0.017 |
| Baganda | 0.271 | 0.325 | 0.193 | 0.040 | 0.007 | 0.026 | 0.000 | 0.009 | 0.276 | 0.322 | 0.221 | 0.023 | 0.000 | 0.004 | 0.000 | 0.001 | 0.008 | 0.069 | 0.000 | 0.017 | 0.051 | 0.082 | 0.023 | 0.019 |
| Zimbabwean | 0.480 | 0.530 | 0.430 | 0.034 | 0.000 | 0.001 | 0.000 | 0.000 | 0.330 | 0.367 | 0.285 | 0.031 | 0.000 | 0.000 | 0.000 | 0.000 | 0.000 | 0.000 | 0.000 | 0.000 | 0.066 | 0.093 | 0.026 | 0.020 |
| Mozambican | 0.580 | 0.744 | 0.489 | 0.080 | 0.002 | 0.010 | 0.000 | 0.003 | 0.271 | 0.348 | 0.206 | 0.037 | 0.001 | 0.007 | 0.000 | 0.002 | 0.000 | 0.000 | 0.000 | 0.000 | 0.049 | 0.082 | 0.011 | 0.024 |

| ADMIXTURE components | 'GIH' | | | | 'Naro' | | | | 'Kalenjin' | | | | "BaPhuthi':'Naro' | | | | Sum of 'BaPhuthi' &'Naro' | | | |
|---|---|---|---|---|---|---|---|---|---|---|---|---|---|---|---|---|---|---|---|---|
| Population | μ | Max | Min | σ | μ | Max | Min | σ | μ | Max | Min | σ | μ | Max | Min | σ | μ | Max | Min | σ |
| !Xun | 0.000 | 0.004 | 0.000 | 0.001 | 0.771 | 0.886 | 0.632 | 0.068 | 0.021 | 0.067 | 0.000 | 0.022 | 0.00E+00 | 0.00E+00 | 0.00E+00 | 0.07 | 0.77 | 0.89 | 0.63 | 0.00E+00 |
| CDX | 0.000 | 0.000 | 0.000 | 0.000 | 0.000 | 0.000 | 0.000 | 0.000 | 0.000 | 0.000 | 0.000 | 0.000 | 0.00E+00 | 0.00E+00 | 0.00E+00 | 0.00 | 0.00 | 0.00 | 0.00 | 0.00E+00 |
| CHB | 0.002 | 0.025 | 0.000 | 0.006 | 0.000 | 0.000 | 0.000 | 0.000 | 0.000 | 0.000 | 0.000 | 0.000 | 0.00E+00 | 0.00E+00 | 0.00E+00 | 0.00 | 0.00 | 0.00 | 0.00 | 0.00E+00 |
| FIN | 0.002 | 0.028 | 0.000 | 0.006 | 0.000 | 0.001 | 0.000 | 0.000 | 0.001 | 0.009 | 0.000 | 0.002 | 0.00E+00 | 0.00E+00 | 0.00E+00 | 0.00 | 0.00 | 0.00 | 0.00 | 0.00E+00 |
| IBS | 0.000 | 0.007 | 0.000 | 0.002 | 0.000 | 0.002 | 0.000 | 0.001 | 0.000 | 0.001 | 0.000 | 0.000 | 0.00E+00 | 0.00E+00 | 0.00E+00 | 0.00 | 0.00 | 0.00 | 0.00 | 0.00E+00 |
| KHV | 0.009 | 0.030 | 0.000 | 0.011 | 0.000 | 0.004 | 0.000 | 0.001 | 0.000 | 0.000 | 0.000 | 0.000 | 0.00E+00 | 0.00E+00 | 0.00E+00 | 0.00 | 0.00 | 0.00 | 0.00 | 0.00E+00 |
| Naro | 0.000 | 0.000 | 0.000 | 0.000 | 0.982 | 0.989 | 0.975 | 0.010 | 0.000 | 0.000 | 0.000 | 0.000 | 0.00E+00 | 0.00E+00 | 0.00E+00 | 0.01 | 0.98 | 0.99 | 0.98 | 0.00E+00 |
| TSI | 0.019 | 0.038 | 0.000 | 0.011 | 0.000 | 0.000 | 0.000 | 0.000 | 0.000 | 0.000 | 0.000 | 0.000 | 0.00E+00 | 0.00E+00 | 0.00E+00 | 0.00 | 0.00 | 0.00 | 0.00 | 0.00E+00 |
| Ju\|'hoansi | 0.000 | 0.000 | 0.000 | 0.000 | 0.967 | 1.000 | 0.534 | 0.124 | 0.006 | 0.077 | 0.000 | 0.021 | 9.76E-03 | 1.37E-01 | 0.00E+00 | 0.10 | 0.97 | 1.00 | 0.61 | 3.65E-02 |

| | | | | | | | | | | | | | | | | | | | | |
|---|---|---|---|---|---|---|---|---|---|---|---|---|---|---|---|---|---|---|---|---|
| Hai\|\|om | 0.003 | 0.012 | 0.000 | 0.005 | 0.501 | 0.519 | 0.488 | 0.011 | 0.047 | 0.066 | 0.016 | 0.018 | 3.77E-02 | 1.14E-01 | 0.00E+00 | 0.03 | 0.52 | 0.56 | 0.49 | 4.76E-02 |
| Nama | 0.009 | 0.035 | 0.000 | 0.011 | 0.589 | 0.857 | 0.150 | 0.208 | 0.006 | 0.071 | 0.000 | 0.017 | 7.33E-02 | 3.85E-01 | 0.00E+00 | 0.20 | 0.62 | 0.86 | 0.21 | 1.03E-01 |
| Jola | 0.000 | 0.000 | 0.000 | 0.000 | 0.006 | 0.017 | 0.000 | 0.005 | 0.002 | 0.034 | 0.000 | 0.008 | 4.20E-02 | 5.62E-01 | 0.00E+00 | 0.01 | 0.01 | 0.02 | 0.00 | 1.37E-01 |
| Khwe | 0.001 | 0.004 | 0.000 | 0.001 | 0.359 | 0.438 | 0.226 | 0.056 | 0.023 | 0.073 | 0.000 | 0.027 | 1.33E-01 | 4.59E-01 | 0.00E+00 | 0.04 | 0.40 | 0.47 | 0.33 | 1.42E-01 |
| ‡Khomani | 0.045 | 0.118 | 0.000 | 0.043 | 0.644 | 1.000 | 0.377 | 0.191 | 0.001 | 0.015 | 0.000 | 0.003 | 1.71E-01 | 6.89E-01 | 0.00E+00 | 0.21 | 0.75 | 1.00 | 0.41 | 1.84E-01 |
| Oromo | 0.014 | 0.031 | 0.000 | 0.011 | 0.015 | 0.037 | 0.003 | 0.009 | 0.106 | 0.175 | 0.066 | 0.029 | 7.67E-02 | 7.30E-01 | 0.00E+00 | 0.01 | 0.02 | 0.04 | 0.00 | 1.89E-01 |
| Chrissie San | 0.000 | 0.000 | 0.000 | 0.000 | 0.449 | 0.486 | 0.394 | 0.048 | 0.000 | 0.000 | 0.000 | 0.000 | 1.21E+00 | 1.44E+00 | 1.06E+00 | 0.02 | 0.99 | 1.00 | 0.96 | 1.98E-01 |
| Himba | 0.000 | 0.000 | 0.000 | 0.000 | 0.068 | 0.084 | 0.047 | 0.013 | 0.000 | 0.000 | 0.000 | 0.000 | 7.60E-02 | 6.08E-01 | 0.00E+00 | 0.02 | 0.07 | 0.12 | 0.05 | 2.15E-01 |
| Karretjie | 0.031 | 0.071 | 0.000 | 0.025 | 0.710 | 0.945 | 0.431 | 0.154 | 0.001 | 0.011 | 0.000 | 0.003 | 2.97E-01 | 1.23E+00 | 1.43E-02 | 0.09 | 0.89 | 1.00 | 0.72 | 2.80E-01 |
| G\|ui & G\|\|ana | 0.000 | 0.000 | 0.000 | 0.000 | 0.682 | 0.843 | 0.471 | 0.123 | 0.008 | 0.039 | 0.000 | 0.013 | 4.74E-01 | 1.06E+00 | 1.26E-01 | 0.02 | 0.97 | 1.00 | 0.95 | 3.02E-01 |
| BaPhuthi_SA | 0.000 | 0.000 | 0.000 | 0.000 | 0.152 | 0.166 | 0.138 | 0.020 | 0.023 | 0.046 | 0.000 | 0.032 | 5.09E+00 | 5.47E+00 | 4.71E+00 | 0.04 | 0.92 | 0.95 | 0.89 | 5.34E-01 |
| Mbukushu | 0.001 | 0.005 | 0.000 | 0.002 | 0.062 | 0.076 | 0.054 | 0.008 | 0.065 | 0.108 | 0.033 | 0.024 | 4.07E+00 | 4.91E+00 | 3.17E+00 | 0.02 | 0.31 | 0.34 | 0.29 | 6.11E-01 |
| AmaNdebele | 0.000 | 0.003 | 0.000 | 0.001 | 0.177 | 0.216 | 0.135 | 0.031 | 0.020 | 0.069 | 0.000 | 0.025 | 4.07E+00 | 5.41E+00 | 3.18E+00 | 0.02 | 0.88 | 0.91 | 0.84 | 8.32E-01 |
| BaPhuthi_LE | 0.001 | 0.013 | 0.000 | 0.003 | 0.149 | 0.201 | 0.116 | 0.022 | 0.004 | 0.045 | 0.000 | 0.010 | 5.53E+00 | 6.96E+00 | 3.45E+00 | 0.04 | 0.95 | 1.00 | 0.86 | 9.52E-01 |
| AmaZulu | 0.000 | 0.001 | 0.000 | 0.000 | 0.128 | 0.169 | 0.095 | 0.019 | 0.012 | 0.047 | 0.000 | 0.015 | 6.07E+00 | 8.10E+00 | 4.25E+00 | 0.02 | 0.89 | 0.92 | 0.84 | 9.84E-01 |
| DumaSan | 0.005 | 0.026 | 0.000 | 0.011 | 0.160 | 0.226 | 0.142 | 0.037 | 0.019 | 0.032 | 0.000 | 0.014 | 4.75E+00 | 5.44E+00 | 2.55E+00 | 0.05 | 0.88 | 0.93 | 0.80 | 1.23E+00 |
| Damara | 0.000 | 0.000 | 0.000 | 0.000 | 0.145 | 0.249 | 0.024 | 0.078 | 0.000 | 0.000 | 0.000 | 0.000 | 6.34E-01 | 4.66E+00 | 0.00E+00 | 0.07 | 0.17 | 0.25 | 0.07 | 1.63E+00 |
| BaSotho | 0.000 | 0.000 | 0.000 | 0.000 | 0.143 | 0.220 | 0.064 | 0.042 | 0.023 | 0.070 | 0.000 | 0.022 | 5.32E+00 | 1.05E+01 | 3.08E+00 | 0.06 | 0.82 | 0.92 | 0.73 | 2.09E+00 |
| Amhara | 0.011 | 0.028 | 0.000 | 0.009 | 0.006 | 0.019 | 0.000 | 0.006 | 0.072 | 0.095 | 0.042 | 0.016 | 9.86E-01 | 1.84E+01 | 0.00E+00 | 0.01 | 0.01 | 0.02 | 0.00 | 4.11E+00 |
| Kwangali | 0.000 | 0.000 | 0.000 | 0.000 | 0.030 | 0.043 | 0.015 | 0.010 | 0.076 | 0.111 | 0.038 | 0.022 | 9.70E+00 | 1.88E+01 | 5.52E+00 | 0.02 | 0.29 | 0.32 | 0.26 | 4.65E+00 |
| Kongo | 0.000 | 0.000 | 0.000 | 0.000 | 0.014 | 0.026 | 0.007 | 0.007 | 0.100 | 0.119 | 0.071 | 0.016 | 2.30E+01 | 3.89E+01 | 1.09E+01 | 0.02 | 0.28 | 0.30 | 0.27 | 1.12E+01 |
| JPT | 0.003 | 0.012 | 0.000 | 0.004 | 0.000 | 0.000 | 0.000 | 0.000 | 0.000 | 0.000 | 0.000 | 0.000 | 1.00E+16 | 2.00E+17 | 0.00E+00 | 0.00 | 0.00 | 0.00 | 0.00 | 4.47E+16 |
| GBR | 0.006 | 0.021 | 0.000 | 0.006 | 0.001 | 0.007 | 0.000 | 0.002 | 0.001 | 0.007 | 0.000 | 0.002 | 6.50E+16 | 9.00E+17 | 0.00E+00 | 0.00 | 0.00 | 0.01 | 0.00 | 2.16E+17 |
| GIH | 0.927 | 1.000 | 0.636 | 0.121 | 0.001 | 0.010 | 0.000 | 0.002 | 0.000 | 0.000 | 0.000 | 0.000 | 3.30E+17 | 6.60E+18 | 0.00E+00 | 0.00 | 0.00 | 0.01 | 0.00 | 1.48E+18 |
| MKK | 0.005 | 0.026 | 0.000 | 0.008 | 0.007 | 0.016 | 0.000 | 0.005 | 0.540 | 0.588 | 0.464 | 0.031 | 7.65E+17 | 1.53E+19 | 0.00E+00 | 0.02 | 0.02 | 0.07 | 0.00 | 3.42E+18 |
| Mandinka | 0.000 | 0.001 | 0.000 | 0.000 | 0.009 | 0.017 | 0.000 | 0.006 | 0.015 | 0.048 | 0.000 | 0.017 | 1.91E+18 | 3.82E+19 | 0.00E+00 | 0.03 | 0.03 | 0.08 | 0.00 | 8.54E+18 |
| Kalenjin | 0.000 | 0.000 | 0.000 | 0.000 | 0.004 | 0.013 | 0.000 | 0.005 | 0.681 | 0.752 | 0.612 | 0.035 | 4.69E+18 | 3.10E+19 | 0.00E+00 | 0.02 | 0.02 | 0.07 | 0.00 | 1.01E+19 |
| Fula | 0.000 | 0.004 | 0.000 | 0.001 | 0.005 | 0.023 | 0.000 | 0.007 | 0.015 | 0.063 | 0.000 | 0.021 | 4.66E+18 | 4.16E+19 | 0.00E+00 | 0.03 | 0.02 | 0.10 | 0.00 | 1.26E+19 |
| Somali | 0.003 | 0.039 | 0.000 | 0.009 | 0.001 | 0.012 | 0.000 | 0.003 | 0.007 | 0.058 | 0.000 | 0.016 | 9.58E+18 | 4.88E+19 | 0.00E+00 | 0.02 | 0.02 | 0.07 | 0.00 | 1.46E+19 |
| YRI | 0.000 | 0.002 | 0.000 | 0.000 | 0.000 | 0.000 | 0.000 | 0.000 | 0.021 | 0.051 | 0.000 | 0.016 | 1.55E+20 | 1.96E+20 | 1.29E+20 | 0.02 | 0.15 | 0.20 | 0.13 | 1.86E+19 |
| Wolof | 0.001 | 0.010 | 0.000 | 0.002 | 0.003 | 0.019 | 0.000 | 0.005 | 0.018 | 0.062 | 0.000 | 0.022 | 1.44E+19 | 8.90E+19 | 0.00E+00 | 0.02 | 0.02 | 0.09 | 0.00 | 2.50E+19 |
| Ga-Adangbe | 0.000 | 0.001 | 0.000 | 0.000 | 0.004 | 0.011 | 0.000 | 0.004 | 0.012 | 0.051 | 0.000 | 0.014 | 3.18E+19 | 1.51E+20 | 7.12E+00 | 0.02 | 0.13 | 0.16 | 0.09 | 5.74E+19 |
| Kikuyu | 0.002 | 0.017 | 0.000 | 0.004 | 0.006 | 0.014 | 0.000 | 0.005 | 0.395 | 0.458 | 0.332 | 0.032 | 2.96E+19 | 2.32E+20 | 1.01E+01 | 0.03 | 0.19 | 0.24 | 0.15 | 7.36E+19 |
| Igbo | 0.000 | 0.004 | 0.000 | 0.001 | 0.000 | 0.004 | 0.000 | 0.001 | 0.035 | 0.095 | 0.000 | 0.024 | 1.39E+20 | 2.05E+20 | 4.23E+01 | 0.02 | 0.17 | 0.20 | 0.13 | 7.44E+19 |

| | | | | | | | | | | | | | | | | | | | |
|---|---|---|---|---|---|---|---|---|---|---|---|---|---|---|---|---|---|---|---|
| LWK | 0.001 | 0.008 | 0.000 | 0.002 | 0.000 | 0.004 | 0.000 | 0.001 | 0.474 | 0.528 | 0.419 | 0.026 | 2.11E+20 | 2.93E+20 | 5.78E+01 | 0.03 | 0.24 | 0.29 | 0.19 | 7.66E+19 |
| Barundi | 0.003 | 0.018 | 0.000 | 0.006 | 0.009 | 0.022 | 0.000 | 0.007 | 0.348 | 0.439 | 0.289 | 0.035 | 4.54E+19 | 2.77E+20 | 8.93E+00 | 0.03 | 0.23 | 0.28 | 0.18 | 9.42E+19 |
| Banyarwanda | 0.002 | 0.011 | 0.000 | 0.003 | 0.007 | 0.022 | 0.000 | 0.008 | 0.365 | 0.434 | 0.323 | 0.031 | 7.88E+19 | 2.58E+20 | 6.76E+00 | 0.04 | 0.20 | 0.26 | 0.11 | 1.11E+20 |
| Angolan | 0.000 | 0.000 | 0.000 | 0.000 | 0.010 | 0.027 | 0.000 | 0.009 | 0.089 | 0.117 | 0.068 | 0.014 | 6.75E+19 | 3.06E+20 | 9.77E+00 | 0.03 | 0.27 | 0.31 | 0.23 | 1.27E+20 |
| Yansi | 0.000 | 0.000 | 0.000 | 0.000 | 0.006 | 0.011 | 0.000 | 0.006 | 0.103 | 0.124 | 0.080 | 0.018 | 6.91E+19 | 2.76E+20 | 2.21E+01 | 0.02 | 0.28 | 0.30 | 0.25 | 1.38E+20 |
| Zambian | 0.000 | 0.000 | 0.000 | 0.000 | 0.011 | 0.046 | 0.000 | 0.014 | 0.126 | 0.179 | 0.090 | 0.028 | 4.72E+19 | 4.25E+20 | 7.98E+00 | 0.04 | 0.38 | 0.42 | 0.33 | 1.42E+20 |
| Baganda | 0.001 | 0.007 | 0.000 | 0.002 | 0.002 | 0.009 | 0.000 | 0.003 | 0.385 | 0.429 | 0.336 | 0.027 | 1.74E+20 | 3.25E+20 | 2.76E+01 | 0.04 | 0.27 | 0.33 | 0.19 | 1.47E+20 |
| Zimbabwean | 0.000 | 0.000 | 0.000 | 0.000 | 0.004 | 0.019 | 0.000 | 0.006 | 0.120 | 0.149 | 0.092 | 0.016 | 3.20E+20 | 5.30E+20 | 2.66E+01 | 0.03 | 0.48 | 0.53 | 0.43 | 2.38E+20 |
| Mozambican | 0.000 | 0.001 | 0.000 | 0.000 | 0.005 | 0.019 | 0.000 | 0.006 | 0.093 | 0.165 | 0.018 | 0.040 | 2.56E+20 | 6.58E+20 | 2.64E+01 | 0.08 | 0.59 | 0.75 | 0.49 | 2.90E+20 |

*Table_S 4: Results of formal test for admixture using $F_3$ estimates.*

*See SuppTable_All_22020X.xlsx file.*

*Table_S 5: Admixture date estimates considering pairs of sources using linkage disequilibrium decay curves. Detected events have been grouped by the language and regions of the identified sources and the date estimates. Where multiple events were detected, the number of events is indicated under 'event'. Symbols : μ – Mean, σ – Standard deviation.*

| Generations ago | | | | | Source pairs | | | | Date (CE) | | |
|---|---|---|---|---|---|---|---|---|---|---|---|
| μ | σ | Z-score | Lower bound | Upper bound | | Population | Language group | | μ | Lower bound | Upper bound |
| 383.28 | 47.15 | 11.0377 | 336.13 | 430.44 | Ju\|'hoansi | CDX | SouthernAfrica_JU | East Eurasian | -8772 | -7452 | -10092 |
| 462.17 | 116.80 | 3.53596 | 345.37 | 578.97 | Himba | CDX | SouthernAfrica_SBW | East Eurasian | -10981 | -7710 | -14251 |
| 239.29 | 25.96 | 10.0017 | 213.33 | 265.25 | Mbukushu | JPT | SouthernAfrica_SBC | East Eurasian | -4740 | -4013 | -5467 |
| 540.40 | 155.02 | 3.03939 | 385.38 | 695.41 | !Xun | GBR | MiddleAfrica_JU | West Eurasian | -13171 | -8831 | -17512 |
| 413.97 | 55.77 | 6.11856 | 358.2 | 469.74 | Ju\|'hoansi | FIN | SouthernAfrica_JU | West Eurasian | -9631 | -8070 | -11193 |
| 429.01 | 104.64 | 4.71743 | 324.37 | 533.66 | Ju\|'hoansi | TSI | SouthernAfrica_JU | West Eurasian | -10052 | -7122 | -12982 |
| 401.19 | 99.14 | 4.88236 | 302.05 | 500.33 | Ju\|'hoansi | TSI | SouthernAfrica_JU | West Eurasian | -9273 | -6497 | -12049 |
| 322.60 | 32.91 | 9.43887 | 289.69 | 355.51 | Ju\|'hoansi | GBR | SouthernAfrica_JU | West Eurasian | -7073 | -6151 | -7994 |
| 327.25 | 63.46 | 8.56548 | 263.79 | 390.71 | Ju\|'hoansi | TSI | SouthernAfrica_JU | West Eurasian | -7203 | -5426 | -8980 |
| 311.83 | 50.75 | 5.78977 | 261.08 | 362.59 | Ju\|'hoansi | GBR | SouthernAfrica_JU | West Eurasian | -6771 | -5350 | -8192 |
| 340.31 | 79.38 | 6.97365 | 260.94 | 419.69 | Ju\|'hoansi | GBR | SouthernAfrica_JU | West Eurasian | -7569 | -5346 | -9791 |
| 299.05 | 41.15 | 9.65892 | 257.9 | 340.2 | Ju\|'hoansi | IBS | SouthernAfrica_JU | West Eurasian | -6413 | -5261 | -7566 |
| 287.10 | 40.93 | 7.06794 | 246.17 | 328.02 | Ju\|'hoansi | TSI | SouthernAfrica_JU | West Eurasian | -6079 | -4933 | -7225 |
| 271.72 | 37.79 | 7.00597 | 233.93 | 309.51 | Ju\|'hoansi | GBR | SouthernAfrica_JU | West Eurasian | -5648 | -4590 | -6706 |
| 198.02 | 34.32 | 15.2452 | 163.71 | 232.34 | Ju\|'hoansi | TSI | SouthernAfrica_JU | West Eurasian | -3585 | -2624 | -4546 |
| 213.19 | 59.30 | 10.4875 | 153.88 | 272.49 | Ju\|'hoansi | TSI | SouthernAfrica_JU | West Eurasian | -4009 | -2349 | -5670 |
| 186.39 | 46.40 | 13.127 | 139.99 | 232.79 | Ju\|'hoansi | GBR | SouthernAfrica_JU | West Eurasian | -3259 | -1960 | -4558 |
| 143.45 | 22.25 | 8.87376 | 121.2 | 165.7 | Ju\|'hoansi | TSI | SouthernAfrica_JU | West Eurasian | -2057 | -1434 | -2679 |
| 105.49 | 4.63 | 34.7354 | 100.86 | 110.11 | Ju\|'hoansi | TSI | SouthernAfrica_JU | West Eurasian | -994 | -864 | -1123 |
| 113.25 | 18.88 | 13.2213 | 94.37 | 132.13 | Ju\|'hoansi | TSI | SouthernAfrica_JU | West Eurasian | -1211 | -682 | -1740 |

| | | | | | | | | | | |
|---|---|---|---|---|---|---|---|---|---|---|
| 80.57 | 4.69 | 27.3558 | 75.88 | 85.25 | Ju\|'hoansi | TSI | SouthernAfrica_JU | West Eurasian | -296 | -165 | -427 |
| 96.46 | 29.03 | 5.1399 | 67.43 | 125.49 | Ju\|'hoansi | TSI | SouthernAfrica_JU | West Eurasian | -741 | 72 | -1554 |
| 72.61 | 8.74 | 15.2149 | 63.87 | 81.34 | Ju\|'hoansi | TSI | SouthernAfrica_JU | West Eurasian | -73 | 172 | -318 |
| 67.98 | 6.99 | 15.3378 | 60.99 | 74.97 | Ju\|'hoansi | TSI | SouthernAfrica_JU | West Eurasian | 57 | 252 | -139 |
| 60.54 | 2.96 | 35.2992 | 57.58 | 63.5 | !Xun | TSI | MiddleAfrica_JU | West Eurasian | 265 | 348 | 182 |
| 46.03 | 2.77 | 29.0446 | 43.26 | 48.8 | Ju\|'hoansi | TSI | SouthernAfrica_JU | West Eurasian | 671 | 749 | 594 |
| 39.06 | 6.36 | 24.9832 | 32.7 | 45.42 | Ju\|'hoansi | TSI | SouthernAfrica_JU | West Eurasian | 866 | 1044 | 688 |
| 44.94 | 13.99 | 3.8417 | 30.95 | 58.93 | Ju\|'hoansi | GBR | SouthernAfrica_JU | West Eurasian | 702 | 1093 | 310 |
| 34.10 | 3.46 | 17.2799 | 30.64 | 37.56 | Ju\|'hoansi | IBS | SouthernAfrica_JU | West Eurasian | 1005 | 1102 | 908 |
| 25.57 | 2.55 | 19.5696 | 23.02 | 28.13 | Ju\|'hoansi | GBR | SouthernAfrica_JU | West Eurasian | 1244 | 1315 | 1172 |
| 22.66 | 3.17 | 6.3083 | 19.5 | 25.83 | Ju\|'hoansi | IBS | SouthernAfrica_JU | West Eurasian | 1325 | 1414 | 1237 |
| 23.12 | 4.91 | 8.3932 | 18.21 | 28.03 | Ju\|'hoansi | IBS | SouthernAfrica_JU | West Eurasian | 1313 | 1450 | 1175 |
| 4.88 | 0.34 | 29.5483 | 4.53 | 5.22 | Ju\|'hoansi | GBR | SouthernAfrica_JU | West Eurasian | 1823 | 1833 | 1814 |
| 5.04 | 0.52 | 19.6198 | 4.52 | 5.56 | Ju\|'hoansi | GBR | SouthernAfrica_JU | West Eurasian | 1819 | 1833 | 1804 |
| 6.20 | 1.69 | 3.65676 | 4.51 | 7.89 | Ju\|'hoansi | TSI | SouthernAfrica_JU | West Eurasian | 1786 | 1834 | 1739 |
| 4.95 | 0.47 | 14.7114 | 4.48 | 5.42 | Ju\|'hoansi | GBR | SouthernAfrica_JU | West Eurasian | 1821 | 1835 | 1808 |
| 27.81 | 8.67 | 3.84146 | 19.14 | 36.48 | Jola | IBS | WesternAfrica_NCA | West Eurasian | 1181 | 1424 | 939 |
| 11.17 | 3.23 | 4.43535 | 7.94 | 14.39 | Jola | IBS | WesternAfrica_NCA | West Eurasian | 1647 | 1738 | 1557 |
| 396.41 | 29.75 | 11.3373 | 366.66 | 426.17 | Yansi | TSI | MiddleAfrica_SBC | West Eurasian | -9140 | -8306 | -9973 |
| 359.22 | 28.68 | 10.9486 | 330.54 | 387.9 | Yansi | TSI | MiddleAfrica_SBC | West Eurasian | -8098 | -7295 | -8901 |
| 303.42 | 43.11 | 6.63558 | 260.31 | 346.53 | Yansi | TSI | MiddleAfrica_SBC | West Eurasian | -6536 | -5329 | -7743 |
| 268.26 | 24.35 | 10.9687 | 243.91 | 292.61 | Yansi | TSI | MiddleAfrica_SBC | West Eurasian | -5551 | -4869 | -6233 |
| 207.27 | 49.88 | 9.17915 | 157.39 | 257.16 | Yansi | IBS | MiddleAfrica_SBC | West Eurasian | -3844 | -2447 | -5240 |
| 171.81 | 29.32 | 7.47332 | 142.49 | 201.13 | Yansi | FIN | MiddleAfrica_SBC | West Eurasian | -2851 | -2030 | -3672 |
| 130.47 | 6.18 | 29.2972 | 124.29 | 136.64 | Kongo | TSI | MiddleAfrica_SBW | West Eurasian | -1693 | -1520 | -1866 |
| 144.00 | 23.06 | 15.8467 | 120.94 | 167.06 | Yansi | TSI | MiddleAfrica_SBC | West Eurasian | -2072 | -1426 | -2718 |
| 66.58 | 5.14 | 21.2212 | 61.44 | 71.72 | Yansi | TSI | MiddleAfrica_SBC | West Eurasian | 96 | 240 | -48 |
| 11.45 | 1.94 | 5.95213 | 9.51 | 13.39 | Yansi | TSI | MiddleAfrica_SBC | West Eurasian | 1639 | 1694 | 1585 |

| | | | | | | | | | | | |
|---|---|---|---|---|---|---|---|---|---|---|---|
| 11.65 | 2.95 | 4.29019 | 8.7 | 14.6 | Yansi | TSI | MiddleAfrica_SBC | West Eurasian | 1634 | 1716 | 1551 |
| 91.70 | 15.89 | 10.7038 | 75.81 | 107.59 | Mozambican | TSI | EasternAfrica_SBE | West Eurasian | -608 | -163 | -1053 |
| 36.28 | 9.08 | 4.78043 | 27.2 | 45.35 | Zambian | IBS | EasternAfrica_SBE | West Eurasian | 944 | 1198 | 690 |
| 29.94 | 6.63 | 3.93431 | 23.31 | 36.57 | Zambian | TSI | EasternAfrica_SBE | West Eurasian | 1122 | 1307 | 936 |
| 22.54 | 4.46 | 5.40709 | 18.08 | 27 | Mozambican | TSI | EasternAfrica_SBE | West Eurasian | 1329 | 1454 | 1204 |
| 13.48 | 3.40 | 4.94577 | 10.08 | 16.87 | Zambian | TSI | EasternAfrica_SBE | West Eurasian | 1583 | 1678 | 1488 |
| 37.80 | 3.21 | 21.8384 | 34.59 | 41 | Ju\|'hoansi | YRI | SouthernAfrica_JU | WesternAfrica_VC | 902 | 991 | 812 |
| 33.80 | 9.28 | 4.64816 | 24.52 | 43.08 | Ju\|'hoansi | YRI | SouthernAfrica_JU | WesternAfrica_VC | 1014 | 1273 | 754 |
| 11.86 | 1.94 | 6.07822 | 9.92 | 13.8 | Ju\|'hoansi | YRI | SouthernAfrica_JU | WesternAfrica_VC | 1628 | 1682 | 1574 |
| 4.19 | 1.02 | 5.14485 | 3.17 | 5.22 | Ju\|'hoansi | YRI | SouthernAfrica_JU | WesternAfrica_VC | 1843 | 1871 | 1814 |
| 49.19 | 6.06 | 11.2199 | 43.14 | 55.25 | Ju\|'hoansi | Damara | SouthernAfrica_JU | SouthernAfrica_KK | 583 | 752 | 413 |
| 57.06 | 17.16 | 3.79992 | 39.9 | 74.21 | Ju\|'hoansi | Himba | SouthernAfrica_JU | SouthernAfrica_SBW | 362 | 843 | -118 |
| 3.14 | 0.66 | 8.79316 | 2.48 | 3.8 | Ju\|'hoansi | Himba | SouthernAfrica_JU | SouthernAfrica_SBW | 1872 | 1891 | 1854 |
| 27.12 | 2.06 | 17.3253 | 25.05 | 29.18 | Ju\|'hoansi | Mozambican | SouthernAfrica_JU | EasternAfrica_SBE | 1201 | 1258 | 1143 |
